# Transcranial ultrasound stimulation modulates neuronal membrane potentials across broad timescales in the awake mammalian brain

**DOI:** 10.1101/2025.06.09.658675

**Authors:** Emma Bortz, Erynne San Antonio, Jack Sherman, Hua-an Tseng, Laura Raiff, Xue Han

**Affiliations:** Department of Biomedical Engineering, Boston University; Department of Pharmacology, Physiology & Biophysics, Boston University

**Keywords:** Ultrasound, Neuromodulation, Voltage imaging, Entrainment, Membrane potential, Plasticity, Cortical neurons

## Abstract

**Background:** Transcranial ultrasound stimulation (TUS) offers noninvasive neuromodulation with high spatial and temporal precision, but its cellular-level effects in the awake brain remain poorly understood.

**Objective:** We investigated how low-intensity TUS modulates membrane voltage dynamics in single cortical neurons in awake mice.

**Methods:** Using the genetically encoded voltage indicator SomArchon, we performed high-speed kilohertz voltage imaging in awake head-fixed mice. TUS was delivered with a 0.35 MHz transducer at 10 or 40 Hz pulse repetition frequency with a 20% duty cycle, at intensities below the estimated threshold for auditory brainstem activation. We analyzed changes in membrane potential (Vm), spiking, and coordination across simultaneously recorded neurons.

**Results:** TUS evoked rapid (<10 ms) Vm depolarizations in 42.8% of neurons, while only 20.5% showed increased spiking, highlighting a direct effect of TUS on modulating synaptic inputs. Many neurons were entrained at both PRFs (20.8% at 10 Hz; 12.7% at 40 Hz) with Vm exhibiting significant phase-locking to individual TUS pulses. Vm entrainment was accompanied by increased temporal coordination across neurons and reset network synchrony. Furthermore, TUS-evoked cellular responses adapted over time, often transitioning from membrane depolarization to hyperpolarization upon repeated exposures, demonstrating prominent response depression.

**Conclusion:** By resolving single-neuron responses, our results demonstrate that TUS directly activates individual cortical neuron with a latency shorter than 10 ms. TUS pulsed at physiologically relevant frequencies of 10 and 40 Hz robustly entrains neural dynamics, alters network coordination and evokes neuronal plasticity. These results highlight the therapeutic potential of designing TUS pulsing patterns to target specific neural circuit dynamics.

## Introduction

Low-intensity, low-frequency transcranial ultrasound stimulation (TUS) is an emerging noninvasive neuromodulation technique capable of targeting deep brain regions with high spatiotemporal precision. Ultrasound stimulation elicits neural responses in both the central^1–5^ and peripheral nervous systems^6–10^. Several biological and physical factors have been proposed to contribute to TUS-induced changes in neural activity. While the precise physiological mechanisms of low-intensity TUS remain incompletely understood, acoustic radiation force is considered a predominant factor^11,12^, generating submicron membrane displacements that activate mechanosensitive ion channels^13–18^. Other proposed mechanisms include intramembrane cavitation^19–21^ and synaptic vesicle release^22^.

In clinical studies, TUS is often delivered at pulse repetition frequencies (PRFs) of 5–100 Hz^23–28^, primarily to minimize thermal effects and ensure patient safety. However, stimulation frequency is a key factor influencing the therapeutic efficacy of neuromodulation techniques. For example, deep brain stimulation (DBS), commonly used to manage Parkinson’s disease and epilepsy, is typically delivered at high frequencies of 130–300 Hz, as lower frequencies yield inconsistent results^29,30^. In contrast, transcranial magnetic stimulation, primarily used for depression, employs lower frequencies <50 Hz^31^ or bursts of 30-50 Hz pulses at ∼5 Hz (theta-burst protocols)^32–34^. Although the underlying mechanisms remain unclear, lower frequencies (<100 Hz) are thought to promote neural entrainment and better engage intrinsic neural dynamics, while higher frequencies (>100 Hz) may disrupt pathological activity patterns without entrainment^35,36^. Supporting this, our recent voltage imaging study showed that 40 Hz DBS robustly entrained neurons in awake mice, whereas 140 Hz induced a robust informational lesion without prominent entrainment^37^.

TUS serves as an external input to neurons, with its effects shaped by the intrinsic properties of the targeted neurons^38^. TUS-evoked responses can be amplified by voltage-gated ion channels^22,39^; notably, TUS can also elevate cytosolic calcium via Ca^2+^ channels in brain slices, independent of Na⁺ channel activation^40^. These findings underscore the importance of voltage-gated ion channel kinetics in shaping membrane dynamics, which constrain action potentials to a few hundred hertz^41–43^. Although TUS utilizes fundamental frequencies around 0.5–2 MHz to ensure skull penetration, pulsing TUS at physiologically relevant timescales differentially modulate calcium dynamics^15,40^ and spiking activity^44,45^ in individual neurons. For instance, 40 Hz TUS enhances gamma-band oscillations, improves memory performance, and reduces amyloid pathology in Alzheimer’s disease models^46,47^, mirroring effects observed with 40 Hz visual flickers^48^. Mechanistically, these frequency-specific effects likely reflect interactions between acoustic radiation force and intrinsic membrane properties, including mechanosensitive ion channel kinetics, membrane time constants, and neuronal resonance. While excitatory and inhibitory neurons differ in PRF sensitivity, our recent work suggests that variability in TUS responses emerges more strongly at the single-neuron level than across canonical cell types^15^.

Beyond immediate effects, TUS can induce sustained neuroplasticity. In humans, TUS reduces cortical GABA levels and enhances functional connectivity, consistent with lasting disinhibition^49^. TUS also increases motor cortical excitability for up to 30 minutes post-sonication, an effect abolished by blockade of Na⁺/Ca²⁺ channels, N-methyl-D-aspartate (NMDA) receptors, and GABAergic signaling^50^. Mechanistic studies in rodents show that TUS-induced potentiation of motor responses requires Ca²⁺-activated bestrophin-1 channels, NMDA receptors, brain-derived neurotrophic factor, and tropomyosin receptor kinase B^51^. In hippocampal circuits, TUS directed at the medial perforant pathway induces long-term depression of field excitatory postsynaptic potentials, similar to low-frequency electrical stimulation^52^. Thus, TUS engages both rapid neuronal activity changes and slower plasticity-mediated processes to modulate circuit dynamics.

TUS consistently modulates brain activity across diverse experimental models, influencing behavioral changes^53,54^; hemodynamic responses^55,56^; population-level local field potentials^57,58^; cytosolic calcium concentrations^15,59–61;^ and individual neuron spiking patterns^44,45^. However, these studies lack the temporal resolution to capture rapid dynamics evoked by TUS, due to either the intrinsic slowness of the signals, e.g., hemodynamics and cytosolic calcium, or the sparseness of the signals, e.g., spiking that requires temporal integration to detect rate changes. Thus far, the reported latencies of TUS-evoked neural responses range from ∼50–500 ms^14,44,59^, insufficient to confirm a direct neuromodulatory effect. Compounding this complexity is the possibility that TUS does not solely act through direct modulation of neuronal Vm. Astrocyte activation by ultrasound can modulate synaptic function via astrocytic glutamate release^62^. Moreover, TUS, when strong enough, can stimulate the auditory system, subsequently triggering cortical calcium influx and motor twitches hundreds of milliseconds later^63,64^. However, using ramped or lower-intensity waveforms can mitigate these auditory confounds^65,66^. Nonetheless, robust TUS effects persist in deafened mice^45,66,67^, invertebrates^68–70^, and cell cultures that lack auditory inputs^14,71,72^.

To evaluate how TUS modulates Vm, which captures synaptic inputs that dictate spiking outputs at the single-neuron level in the mammalian brain, we performed cellular voltage imaging in the cortex of awake mice using the genetically encoded voltage indicator SomArchon^73^. We delivered low TUS at PRFs of 10 and 40 Hz, known to engage cortical neurons^74–76^, at a low-intensity estimated to not evoke a auditory brainstem response^65^, and characterized the effects of TUS on Vm amplitude, entrainment, spiking, and plasticity.

## Materials and Methods

### Animal Preparation

All procedures were approved by the Boston University Institutional Animal Care and Use Committee (IACUC) and Biosafety Committee. Mice were group-housed prior to surgery and individually housed postoperatively with environmental enrichment, such as igloos or running wheels. The animal facility maintained a 12-hour light/dark cycle at approximately 70 °F and 50% humidity. The study included a total of 20 adult mice, aged 8-16 weeks at experiment onset, including 8 C57BL/6 (Jackson Laboratory #000664, 3 female, 5 male), 5 NDNF-IRES-Cre transgenic mice (Jackson Laboratory #030757, 2 female, 3 male), 5 PV-tdTomato transgenic mice (1 male, 4 female) obtained by crossing B6.Cg-Gt(ROSA)26Sortm14(CAG-tdTomato)Hze/J (Jackson Laboratory, #007914) with B6.129P2-Pvalb^tm1(cre)Arbr^/J (Jackson Laboratory, #017320) mice, and 2 PV-Cre (B6.129P2-Pvalbtm1(cre)Arbr/J, Jackson Laboratory #017320, 2 male).

Surgical preparations were as previous described^37,59^. Briefly, mice were anesthetized with 1–3% isoflurane, and a 3 mm craniotomy was made over either the motor cortex (16 mice; AP: +1.75 mm, ML: 1.75 mm) or visual cortex (4 mice; AP: −2.8 mm, ML: 2.5 mm). AAV vectors were infused through a 36-gauge stainless steel needle (World Precision Instruments, NF36BL-2) connected to a 10 µL NANOFIL microsyringe (World Precision Instruments) and controlled by an UltraMicroPump3-4 microinjector (World Precision Instruments). Injections were made at 2–6 sites per craniotomy, terminating 180–250 µm below the dura. For NDNF-Cre mice, a total of 300 nL of AAV9-Syn-FLEX-SomArchon-GFP (titer: 1.28×10^12^GC/mL) was infused at 50 nL/minute. C57BL/6 mice and the PV-Cre mouse received a total 400 nL of AAV8-CaMKII-SomArchon-GFP (titer: 3.2×10^12^ GC/mL), and PV-tdT mice and sham mice received 400 nL of AAV9-Synapsin-SomArchon-GFP (titer: 5.42×10^12^ GC/mL). After each injection, the needle was left in place for 5–10 min to aid viral diffusion. A #0 glass coverslip (3 mm outer diameter, Warner Instruments, 64-0726) was placed over the craniotomy and sealed to the skull using either surgical silicone adhesive (Kwik-Sil, World Precision Instruments) or ultraviolet-curable dental cement (Tetric EvoFlow, Ivoclar). Exposed skull regions were reinforced with Metabond Quick Adhesive Cement (Parkell, S380), and a custom aluminum headbar was affixed with dental cement (Stoelting, 5145).

All mice received 72 hours of postoperative analgesia via a single preoperative intramuscular injection of sustained-release buprenorphine (0.03 mg/kg; Reckitt Benckiser Healthcare). Mice were allowed to recover for 3–4 weeks before imaging.

### High Speed Voltage Imaging

Mice were habituated to head fixation on the imaging platform for five consecutive days prior to voltage recordings. During each session, neurons expressing SomArchon were identified by detecting GFP fluorescence, which is co-expressed with SomArchon. Voltage imaging was conducted either using a custom widefield fluorescence microscope as described previously^73,37^, or, most of the time, a targeted illumination confocal microscope (TICO) as described previously^97^. Briefly, the widefield microscope was equipped with a Hamamatsu ORCA Fusion Digital sCMOS camera (Hamamatsu Photonics K.K., C14440-20UP), a 40x NA = 0.8 water immersion objective (Nikon, CFI APO NIR), a 140-mW 637 nm laser (Coherent Obis 637-140X) coupled with a reverse 2x beam expander (ThorLabs Inc., GBE02-E) to achieve an illumination with a diameter of 30–40 µm, and a mechanical shutter (Newport corp., model 76995) controlled laser timing via a NI DAQ board (USB-6259). Imaging frames were captured using HCImage Live (Hamamatsu Photonics K.K., U11158) at ∼828 Hz (16 bits, 2 × 2 binning), and data were stored as DCAM image files (DCIMG). The TICO microscope was equipped with a 637 nm diode laser (Ushio America Inc., Red-HP-63X) for SomArchon excitation, a 488 nm laser (Lasertack, GmbH277 PD-01376) for GFP excitation, an sCMOS camera (Teledyne Photometrix, Model: Kinetix), DMD module (Vialux, V-7000 VIS) for targeted illumination, and an 16X objective (Nikon Corp., 16×/0.8NA LWD). Image frames were captured using Teledyne Photometrics PVCAM software (Teledyne Photometrics, PVCAM).

### Transcranial Ultrasound Stimulation

A planar ultrasound transducer with a center frequency of 350 kHz (Ultran GS350-D13) was positioned beneath the chin of awake, head-fixed mice, aligned under the microscope objective. Ultrasound gel (Aquasonic Clear® Ultrasound Gel 03-08_BX, Parker Laboratories, Fairfield, NJ) was applied between the transducer and the chin to ensure efficient acoustic coupling and propagation to the brain. Ultrasound was delivered in 1-second bursts per trial, pulsed at either 10 or 40 Hz with a 20% duty cycle. Pulse trains were triggered via MATLAB (MathWorks, Inc.) through a USB-6259 NI-DAQ multifunction I/O system (National Instruments). TTL triggers from the NI-DAQ and the imaging camera were simultaneously recorded using the Open Ephys platform (http://open-ephys.org) for precise offline alignment of stimulation and imaging frames. Sham stimulation controls were performed by coupling the transducer, with ultrasound gel, to the ventral abdomen of awake, head-fixed mice.

### Acoustic Field Characterization and Simulation

Free-field acoustic pressure was measured in degassed, deionized water using a calibrated 0.2 mm needle hydrophone (Precision Acoustics, UK). The hydrophone was mounted in a custom-built 3D motorized scanning tank constructed according to an open-source design^98^. Pressure waveforms were acquired with a digital oscilloscope (TDS2024C, Tektronix) and processed in MATLAB to extract peak acoustic pressures across a 3D grid centered on the transducer’s focus. Numerical simulations were performed using the k-Wave MATLAB toolbox to model ultrasound propagation through a C57BL/6 mouse skull^99^. The simulation domain incorporated anatomically realistic geometry and acoustic properties of the skull, brain, and surrounding soft tissues, including density, sound speed, and attenuation coefficients derived from the literature, as previously detailed^15,59^. To model realistic acoustic heterogeneity, a static air-filled cavity was incorporated along the nasal cavity and through the pharyngeal airway. The air cavity was modeled as a rectangular prism ∼1 mm^2^ in cross-section, with a gradual dorsoventral taper extending from the start to the end of the pharyngeal airway. The ultrasound source was modeled as a 13 mm diameter circular array. Peak intracranial pressures and spatial peak pulse average intensities (I_SPPA_) were computed directly from k-Wave outputs at each voxel in the simulated brain volume. Spatial peak temporal average intensity (I_SPTA_) was computed by multiplying I_SPPA_ by the pulse duty cycle (20%).

### SomArchon Fluorescence Trace Preprocessing

SomArchon fluorescence traces were analyzed in MATLAB (MathWorks, R2021a). Image frames were first motion-corrected on a per-trial basis, following procedures described previously^37^. Briefly, each trial video was boxcar smoothed over 21 frames and sharpened using a Gaussian filter (filter size = 50; filter sigma = 1 and 25, respectively). The processed videos were then input into the NormCorre algorithm to estimate X and Y frame shifts, using a bin width of 10, maximum shift of 50 pixels, and no upsampling^100^. Each trial was individually motion-corrected. After within-trial correction, the mean motion-corrected images from all trials were aligned across trials using NormCorre. The resulting X and Y shifts from the cross-trial alignment were applied to each frame of the respective trials to generate fully motion-corrected videos.

Following motion correction, regions of interest (ROIs) corresponding to individual neurons were manually selected using the drawpolygon tool in MATLAB. For each ROI, the SomArchon fluorescence trace was computed by averaging the pixel intensities within the selected region. Traces were then interpolated to 1000 Hz using piecewise cubic Hermite interpolation (pchip) to facilitate alignment with ultrasound stimulation timestamps. To remove slow baseline fluctuations, traces were detrended by subtracting a boxcar-smoothed version of the signal using a sliding 1-second window.

Detrended traces were visually inspected, and those containing obvious motion artifacts or lacking discernible spiking activity were excluded from further analysis. The remaining detrended traces, referred to as membrane potential (Vm) traces, were used in all subsequent analyses.

### Spike Identification

Spikes were detected in the Vm traces using a custom algorithm as detailed previously^101^. To estimate spontaneous Vm fluctuations, traces were first high-pass filtered using a 10 ms moving median filter, and then smoothed using a 201 ms moving average to generate a smoothed trace (ST). An asymmetric separation of sub- and supra-threshold fluctuations was applied. The lower trace (*TL*) was computed by replacing all Vm values greater than ST with ST values, as defined in Equation 1:

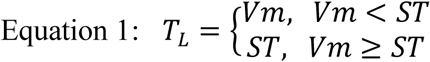

To estimate periods with spiking, we computed the upper trace (*TU*) by replacing all values of the Vm trace below the ST with the ST using Equation 2:

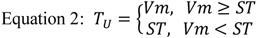

Next, we computed the rate changes in *T_L_* and *T_U_* by calculating their derivatives (*dT_L_* and *dT_U_*) respectively. We then identify putative spikes as the timepoints where *dT_U_* was greater than mean (*μ*) *dT_U_* signal plus 6x the standard deviation (*σ*) of *dT_L_* using Equation 3:

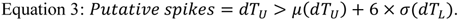

Putative spikes were then evaluated using the following for three criteria. First, putative spikes within the first three and last three datapoints were excluded. Second, the Vm value of the putative spike was greater than the previous point. Third, the datapoint values after the spike drop sharply (i.e., have a negative derivative). That is, *dT*_*U*_ values within the three datapoints after the identified putative spikes were more negative than 5x *σ*(*dT*_*L*_) below the mean. After excluding spikes not meeting the criteria, remaining putative spikes were deemed as final spikes for further analyses.

### Classification of Spike-Modulated and Vm-Modulated Neurons

To determine whether TUS evoked significant changes in a neuron’s Vm, we analyzed two distinct time windows following TUS onset: a transient period (0–250 ms) and a delayed period (250 ms–1 second). During the transient period, the mean Vm within the first 250 ms after TUS onset was calculated for each trial. The mean Vm was then compared to the corresponding pre-stimulation mean Vm of the same trial (1-second pre-TUS onset) using a Wilcoxon signed-rank test (α = 0.05). Neurons with transient mean Vm significantly different than the pre-stimulation mean Vm were further classified as transiently activated or transiently suppressed. For the delayed period (250 ms–1 second), the window was divided into three consecutive 250 ms bins: 250–500 ms, 500–750 ms, and 750–1000 ms. For each bin, the mean Vm was calculated and compared to the pre-stimulation mean Vm for each trial using a Wilcoxon signed-rank test (α = 0.05). A neuron was classified as having delayed activation if at least two consecutive bins showed significant increases in mean Vm compared to the pre-stimulation mean Vm. To avoid overlap between transient and sustained classifications, neurons classified as transiently modulated within the first 250 ms were excluded from the sustained modulation category. Neurons with no significant changes in either time window were classified as non-modulated.

To assess TUS-induced changes in spiking activity, we followed a similar approach. Due to the inherently high variability of spike rates during brief one-second windows, suppression of spike rate during either the transient or the sustained period was not analyzed.

To validate the statistical method used to detect TUS-induced modulation of Vm and spiking activity, we calculated the false positive rate by measuring the percent of false positive neurons when repeating the statistical tests at twenty randomized pseudo-stimulation onsets within the pre-stimulation period. False positive rate for all modulation tests was below 5%, confirming that our statistical approach is robust with a false positive rate below the expected 5% threshold (α = 0.05).

### TUS-evoked Response Latency and Temporal Profiles

Vm latency was analyze across neurons that were classified as Vm transiently activated. Vm traces were first z-scored to the mean of the pre-stimulation period. A Wilcoxon rank-sum test (*p* < 0.05) was then used to compare the instantaneous z-scored Vm for each millisecond time bin across all trials to the mean of the pre-stimulation period (zeros). The first occurrence of a significant point was recorded as Vm latency for a given neuron.

To characterize the temporal dynamics of TUS-evoked responses, we computed the peak time and full-width at half-maximum (FWHM) from trial-averaged Vm and spike rate traces. For each neuron, we identified the time of peak response within a 500 ms stimulation window and defined peak time as the time from TUS onset to this maximum. FWHM was computed as the duration over which the response remained above half the peak amplitude. This was restricted to neurons with significant transient activation.

### Analysis of Evoked Vm Responses Across Repeated TUS Trials

To quantify the change in evoked Vm responses across repeated trials, we z-scored the Vm for each trial relative to the 1-second pre-stimulation period. For each neuron and trial, the response was computed as the mean z-scored Vm within a specific analysis window during stimulation: either the transient period (0–250 ms) or the delayed period (250–1000 ms). The rate of change across trials was then estimated as the slope of the linear regression of these trial-wise mean responses. To determine whether a neuron exhibited a significant change, we applied a threshold based on the slope’s p-value from the linear fit (α < 0.05).

### Vm Phase-Locking Analysis

To evaluate TUS-induced entrainment of Vm, we calculated the phase-locking value (PLV), which measures the consistency of Vm phase relative to stimulation pulse onsets. The PLV was computed with equation 4:

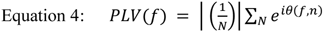

where *f* is the Vm frequency and *N* is the total number of stimulation pulses. The phase *ϕ* of the Vm at the pulse time was obtained from the Hilbert Transform of the narrowband (±5% of *f*) filtered Vm. The average PLV value of each neuron was calculated as the mean across all pulses during all trials. To build the pre-stimulation shuffled distribution for statistical comparison, we randomly shifted the TUS onsets, maintaining pulse repetition frequency, and calculated the mean phase locking value for each iteration. This process was iterated 1500x. If a neuron had a PLV > 95^th^ percentile of the shuffled distribution, the neuron was considered significantly entrained by TUS.

### Vm and Spike Cross-Correlation Between Simultaneously Recorded Neuron Pairs

Pairwise cross-correlation of Vm was computed for each neuron pair within the same field of view (FOV) during the pre-stimulation, stimulation, and post-stimulation periods, then averaged to get one value per FOV. The Vm signal of each neuron was pre-stimulation z-scored for each trial using the mean and standard deviation from the pre-stimulation. Neuron pairwise cross-correlation values were calculated for each trial and then averaged across trials using the MATLAB xcorr function, with zero time lag. Friedman’s test with Wilcoxon signed-rank post-hoc testing was used to assess the significance of changes in correlation coeffects across the pre-stimulation, stimulation, post-stimulation time windows.

Spike cross-correlation was computed using peri-stimulation time histograms (PSTHs) derived from the spike rasters. Spike events were binned at 50 ms intervals (corresponding to 20 Hz effective resolution) and converted into spike rates by dividing by the bin width. PSTHs were trial-averaged and z-scored based on the pre-stimulation period. Pairwise cross-correlation of PSTHs between neuron pairs was calculated at zero lag using the MATLAB xcorr function, following the same procedure as for Vm. Correlation values were averaged across trials, and changes in PSTH cross-correlation across the pre-stimulation, stimulation, and post-stimulation periods were assessed using Friedman’s test with Wilcoxon signed-rank post-hoc test.

### Statistical Analysis

All statistical analyses were performed in MATLAB 2021a using standard non-parametric tests unless otherwise specified. Because data distributions were non-normal, only non-parametric tests (e.g., Wilcoxon signed-rank, rank-sum, Friedman) were used throughout. Statistical test details, including sample sizes and p-values, are reported in the text or summarized in Suppl. Table 3. For circular data, Rayleigh’s test was used. A significance level of α = 0.05 was applied to all tests.

## Results

### Temporally resolved membrane potential analysis of TUS-evoked responses in individual cortical neurons in awake mice

To characterize rapid, real-time neuronal responses evoked by TUS, we performed high-speed voltage imaging in awake, head-fixed mice (**Fig. 1a**). SomArchon^73^, a genetically encoded voltage indicator, was virally expressed in various cortical neuron subtypes (**Fig. 1b, Methods**). A glass coverslip was then placed over the pia to provide optical access. Imaging was conducted using a custom targeted-illumination confocal microscope operating at 828 Hz^77^, allowing simultaneous recording of multiple neurons while avoiding TUS-induced artifacts common to metal electrodes.

**Figure 1:**
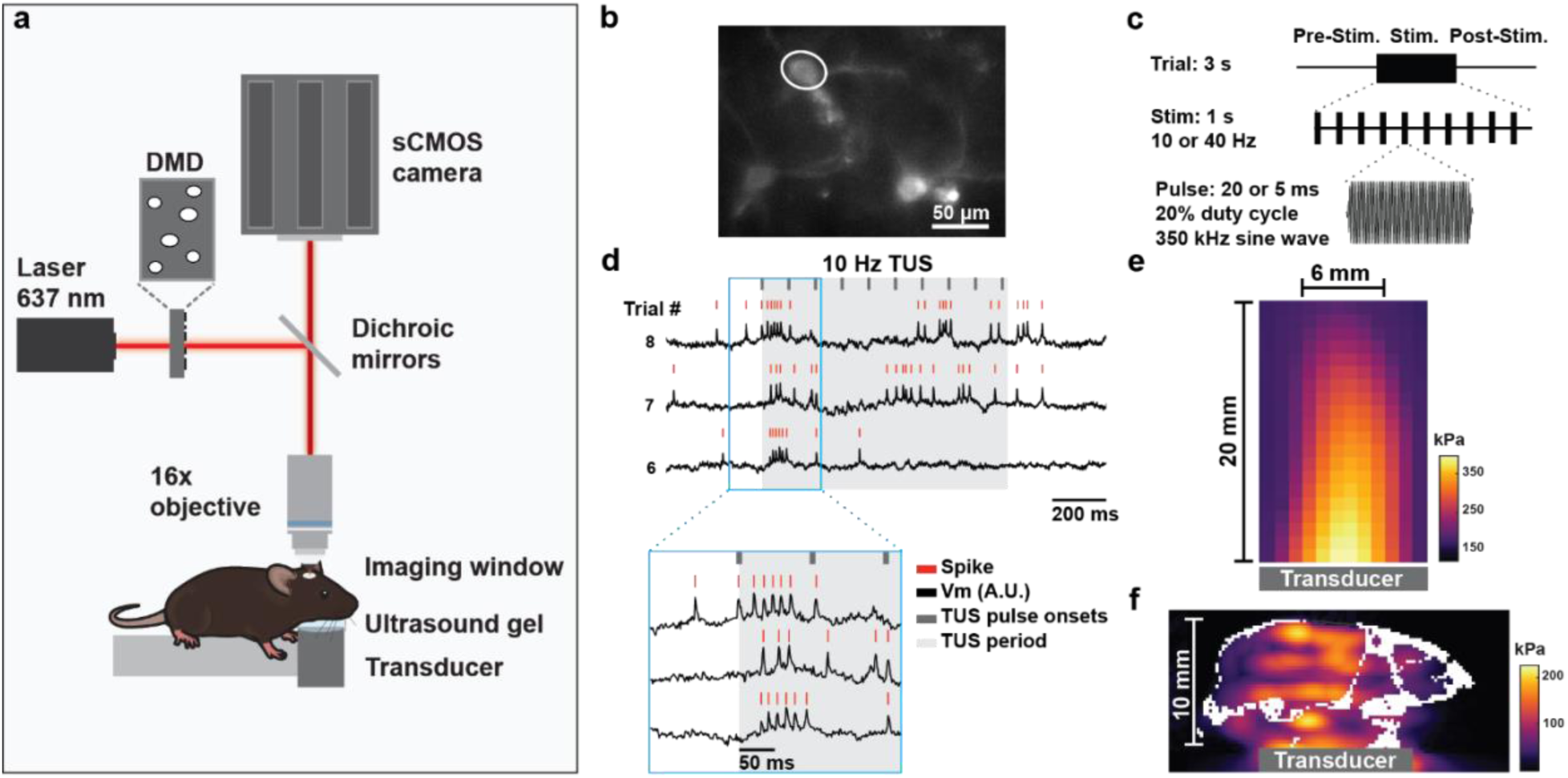
Voltage imaging of cortical neurons during TUS in awake mice. **(a)** Experimental setup: awake mice expressing SomArchon in the cortex were imaged using a custom confocal microscope, equipped with a 637 nm laser, a digital micromirror device (DMD) for patterned excitation, a high-speed sCMOS camera, and a 16× objective. A 6 mm planar transducer delivered TUS via ultrasound gel. **(b)** Example field of view with SomArchon-GFP neurons; Vm and spikes from circled neuron shown in (d) (scale bar: 50 μm). **(c)** TUS protocol: 350 kHz ultrasound, 1 s duration, 20% duty cycle, delivered at 10 or 40 Hz PRF. **(d)** Vm traces of example neurons (circled in (b)) during three trials of 10 Hz TUS. Red ticks: spikes; black: Vm; gray ticks: pulse onsets; shaded gray: 1 s TUS. **(e)** Free-field acoustic pressure in water (peak: 401 kPa). **(f)** Simulated in situ acoustic pressure map in mouse brain using k-Wave (peak: 209 kPa).

TUS was delivered using a 0.35 MHz planar transducer coupled beneath the mouse’s chin with ultrasound gel, ensuring unobstructed optical access (**Fig. 1a**). The planar transducer was selected to match the mouse brain’s dimensions (∼13 × 14 × 8 mm) to the ultrasound focus (∼6.5 × 6.5 × 20 mm at -6 dB). During each 3-second imaging trial, 1 second ultrasound was delivered at either 10 or 40 Hz PRF with a 20% duty cycle (**Fig. 1c; Supplemental Table 1**). Each neuron was typically recorded over 20 trials (range: 10–40), with an inter-trial interval of ∼2-10 seconds. SomArchon fluorescence was processed offline to obtain Vm and spike timing (**Methods**, Fig. 1b-d).

To estimate the ultrasound intensity at the recording site, we first measured the free-field acoustic waveform of the transducer in degassed water using a calibrated needle hydrophone. The peak pressure was 401 kPa with a focal region of 20 mm height and 6 mm width (**Fig. 1e**). Using this empirically measured pressure as input, we performed k-Wave simulations to model intracranial pressure and intensity distributions. Simulations predicted a max intensity of .18 MPa in the soft tissue at the intracranial bone and .11 MPa at the imaging site, the latter corresponding to a maximum in situ I_SPPA_ of 0.129 W/cm^2^ (**Fig. 1f; Supplemental Table 2; methods**). These peak values at the imaging site are consistent with the low-intensity range reported in prior neuromodulation studies^44,52,78,79^, where effective stimulation pressures and I_SPPA_ values ranged from ∼88–438 kPa and ∼0.079–6.4 W/cm^2^, respectively. The intensity at the intracranial bone is below the previously estimated threshold for auditory brainstem activation by Choi et al.^65^.

### TUS induced heterogeneous Vm responses in individual neurons with many exhibiting rapid Vm depolarization within 10 ms of TUS onset, unaccompanied by increased spiking

TUS evoked robust Vm changes across many neurons upon repeated stimulation at both PRFs, though temporal response profiles varied widely (**Fig. 2**). Many neurons exhibited transient depolarization within ∼250 ms of TUS onset (**Fig. 2a-b,e-f**), typically without spike rate increase (**Fig. 2a,e**), although a small fraction exhibited increased spiking (**Fig. 2b,f).** Some neurons displayed delayed Vm depolarization, marked by gradual Vm shifts persisting throughout stimulation (**Fig. 2c,g**), while a smaller group showed consistent hyperpolarization (**Fig. 2d,h**). Most strikingly, many neurons’ Vm fluctuations were entrained by TUS at either 10 or 40 Hz PRFs, faithfully following pulse timing (**Fig. 2g**). The diverse Vm responses evoked by TUS, including depolarization, hyperpolarization, and entrainment, are largely in agreement with that observed during conventional intracranial electrical stimulation^37^. This heterogeneity also aligns with our previous finding that TUS-evoked calcium signals are largely independent of cell type and PRFs^15^.

**Figure 2:**
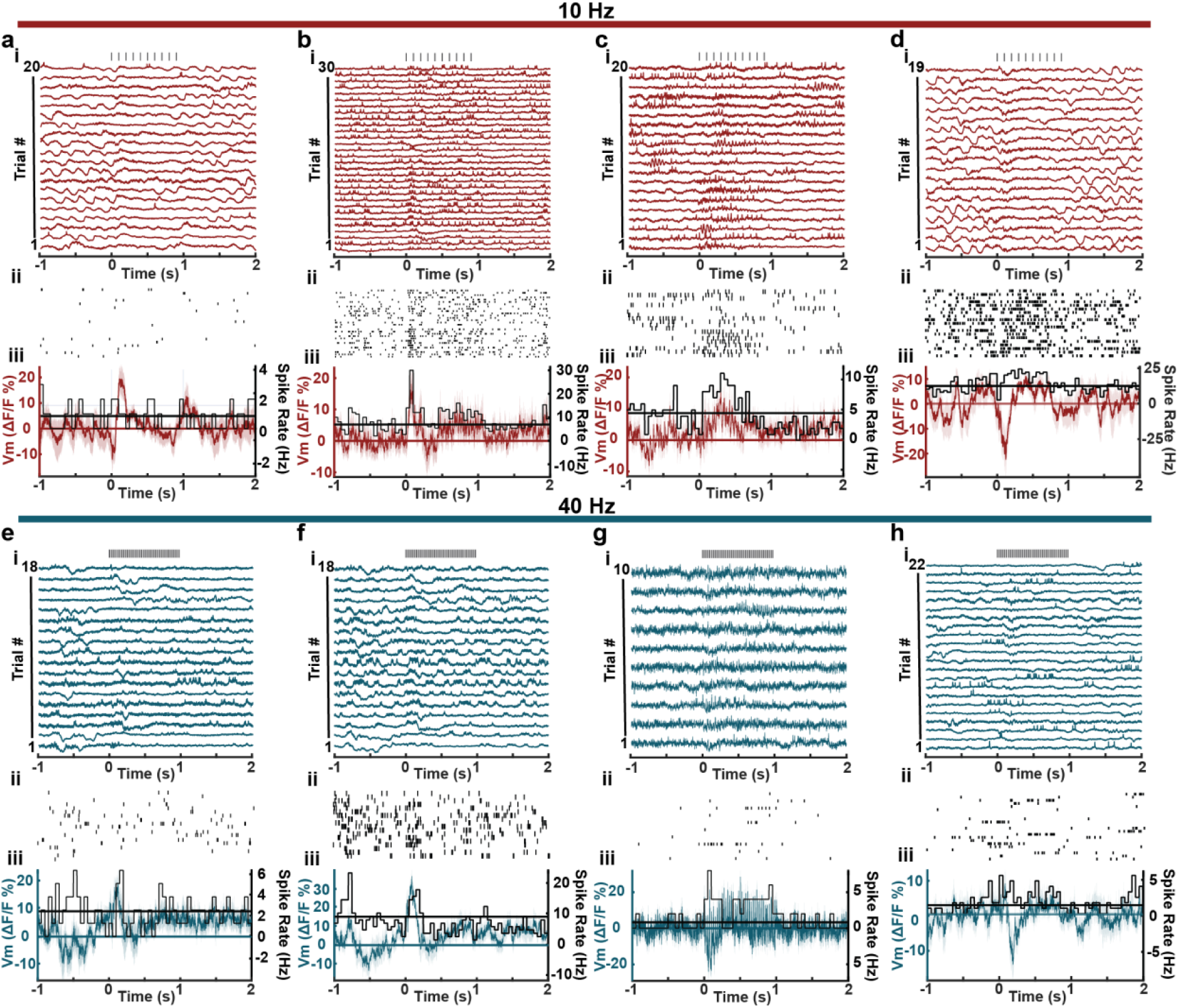
TUS-evoked Vm and spiking responses across example neurons. **(a–d**, red) Example neurons showing TUS-evoked changes in Vm and spike rate during 10 Hz TUS and **(e-h**, blue) during 40 Hz TUS. For each neuron: **(i)** Vm traces across trials, aligned to TUS onset; vertical ticks indicate TUS pulse onsets. **(ii)** Corresponding spike raster plot across trials. **(iii)** Trial-averaged Vm (colored traces, mean ± 95% CI), and spike rate (black histogram, 50 ms bins). Horizontal lines show the average Vm (colored) and spike rate (black) during pre-stimulation, the second before TUS onset.

To assess individual neurons’ Vm responses, we compared mean Vm amplitude during stimulation to pre-stimulation activity, defined as the 1 second preceding TUS onset. Neurons were classified as modulated if Vm significantly exceeded pre-stimulation levels during either the first 250 ms (transient) or 250–1000 ms (delayed) window following stimulation onset (**Methods**, **Fig. 3a-b**). Of 161 recorded neurons across 15 mice, 42.8% (69/161) were significantly modulated, with 34.2% activated (depolarized) and 8.7% suppressed (hyperpolarized) (**Table 1**, **Fig. 3c-f**). Most Vm-modulated neurons (88%, 61/69) responded during the transient window, indicating that rapid Vm shifts dominated TUS responses.

**Figure 3:**
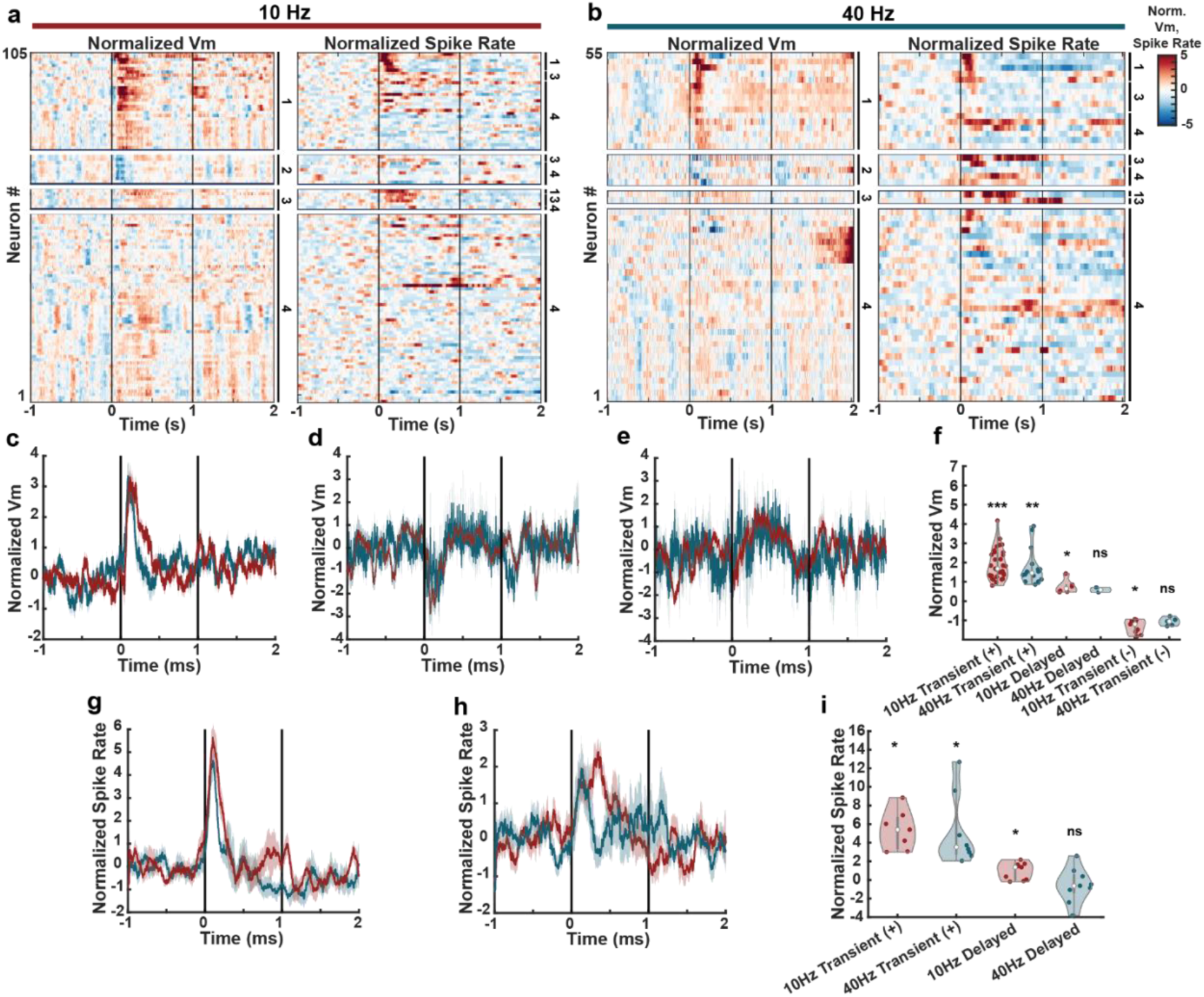
TUS-evoked transient and delayed Vm and spiking changes across individual neurons at 10 and 40 Hz PRFs. **(a–b)** Heatmaps of z-scored Vm (left) and spike rate (right) aligned to TUS onset at 10 Hz **(a**, N = 105) and 40 Hz **(b**, N = 55). Neurons are grouped by response type (1: transient activation, 2: transient suppression, 3: delayed activation, 4: no modulation); spike rate heatmaps are sorted the same as Vm but independently classified. **(c–e)** Mean z-scored Vm for neurons with **(c)** transient activation, **(d)** transient suppression, or **(e)** delayed activation (10 Hz, red: N = 31, 9, 6; 40 Hz, blue: N = 16, 5, 2). **(f)** Violin plots of Vm-modulated neurons (Wilcoxon signed-rank test; see Suppl. Table 3). **(g–h)** Mean z-scored spike rate for neurons with **(g)** transient or **(h)** delayed activation (10 Hz, red: N = 7, 9; 40 Hz, blue: N = 8, 9). **(j)** Violin plots of spike-modulated neurons (Wilcoxon signed-rank test; see Suppl. Table 3).

**Table 1:** Vm and spike modulation across PRFs and conditions.

| TUS |  |  |  |  |  |  |  |
| --- | --- | --- | --- | --- | --- | --- | --- |
| PRF | Metric | Total Neurons | Transient (-) | Transient (+) | Delayed | Total Modulated | Total Modulated (%) |
| 10 | Vm | 106 | 9 | 31 | 6 | 46 | 43.4 |
| 40 | Vm | 55 | 5 | 16 | 2 | 23 | 41.8 |
| **Total** | **Vm** | **161** | **14** | **47** | **8** | **69** | **42.9** |
| 10 | Spike | 106 | - | 7 | 9 | 16 | 15.1 |
| 40 | Spike | 55 | - | 8 | 9 | 17 | 30.1 |
| **Total** | **Spike** | **161** | **-** | **15** | **18** | **33** | **20.5** |

| Sham Ultrasound |  |  |  |  |  |  |  |
| --- | --- | --- | --- | --- | --- | --- | --- |
| 10 | Vm | 39 | 0 | 0 | 0 | 0 | 0.0 |
| 40 | Vm | 21 | 0 | 0 | 0 | 0 | 0.0 |
| **Total** | **Vm** | **60** | **0** | **0** | **0** | **0** | **0.0** |
| 10 | Spike | 39 | - | 0 | 0 | 0 | 0.0 |
| 40 | Spike | 21 | - | 0 | 2 | 2 | 5.1 |
| **Total** | **Spike** | **60** | **-** | **0** | **2** | **2** | **3.3** |

As Vm depolarization often precedes spiking, we next examined spike rate changes during transient and delayed periods (**Methods**). Overall, 20.5% of neurons showed increased spike rates, in agreement with a prior primate study^80^, while none showed significant decreases (**Fig. 3g-i**, **Table 1**). Among spike-modulated neurons, 91% (30/33) also exhibited significant transient Vm depolarization. However, only 45% (15/33) showed transient increases in spiking, compared to 88% of Vm-modulated neurons. Further evaluation revealed that just 23.4% of transiently Vm-activated neurons were spike-modulated, suggesting that many Vm responses occur independently of overt spiking.

As a control, we performed off-target sham ultrasound to the abdomen of the mouse using ultrasound gel. Sham ultrasound induced minimal changes in Vm, with no Vm-modulated neurons (N=60 neurons in 5 mice, **Table 1**). A small fraction of neurons (3.3%, 2/60, **Table 1**) exhibited increased spiking, within the expected 5% false positive rate.

After observing rapid TUS-evoked Vm responses in individual neurons, we further quantified the response latency and the temporal profiles across the transiently Vm-activated neurons (**Methods**). At 10 Hz PRF, TUS evoked rapid depolarizations with a mean latency of 9.58±1.31 ms (range: 5–37 ms, N = 31). Similarly, 40 Hz stimulation produced fast Vm responses with a latency of 13.31±3.94 ms (range: 5–53 ms, N = 16). The slightly longer latency at 40 Hz may reflect reduced energy delivery during the first 20 ms of stimulation, when only a single 5 ms pulse occurred for 40 Hz TUS, compared to a 20 ms pulse for 10 Hz TUS. Despite differences in early energy delivery, the overall activation profiles were broadly comparable across PRFs, with 10 Hz TUS-evoked responses peaking at 154.6±13.3 ms with a full-width at half-maximum (FWHM) of 274.0±23.8 ms and 40 Hz TUS-evoked responses peaking at 117.1±11.7 ms with a FWHM of 199.8±31.9 ms. However, FWHM values were significantly broader at 10 Hz than 40 Hz (Mann–Whitney U test, one-tailed, p = 0.024), indicating more sustained depolarization at 10Hz PRFs. The fact that many neurons exhibited short-latency responses under 10 ms confirms that TUS directly modulates neuronal dynamics at the cellular level, rather than through systemic artifacts, such as heating, or indirect somatosensory or auditory activation.

### TUS-evoked responses systematically change across repeated stimulation trials with more neurons showing response depotentiation

TUS can induce long-lasting changes in neural activity, with effects persisting minutes to hours beyond stimulation offset^81–83^. To assess whether TUS-evoked responses systematically changed across repeated trials, we z-scored the Vm of each trial relative to the 1-second pre-stimulation baseline. Since responses were primarily restricted to the transient period within the first 250 ms after stimulation onset (**Table 1**), our analysis focused on this window.

Many neurons exhibited progressive reductions in TUS-evoked transient Vm responses across trials, often shifting from depolarization to hyperpolarization (**Fig. 4a,c**), while a smaller subset exhibited progressive increases (**Fig. 4b,d**). In total, 19 of 161 neurons (11.8%) showed significant trial-by-trial changes in the transient window (p < 0.05, **Methods**), with 16 progressive decreases and 3 increases. In contrast, during the delayed window (250-1000 ms post-onset) evoked Vm responses showed minimal changes, only 7 neurons (4.3%) showed significant changes across trials, with 6 exhibiting increased Vm and 1 decreased Vm. At the population level, trial-to-trial changes during the transient window were significantly different from zero (Wilcoxon signed-rank test, p = 5.3e-04, **Fig. 4e**), whereas delayed responses were not (p = 0.77, **Fig. 4f**), suggesting a temporally specific effect of evoked response adaptation. Thus, TUS-evoked cellular responses are shaped by plasticity mechanisms and dominated by depotentiation.

**Figure 4:**
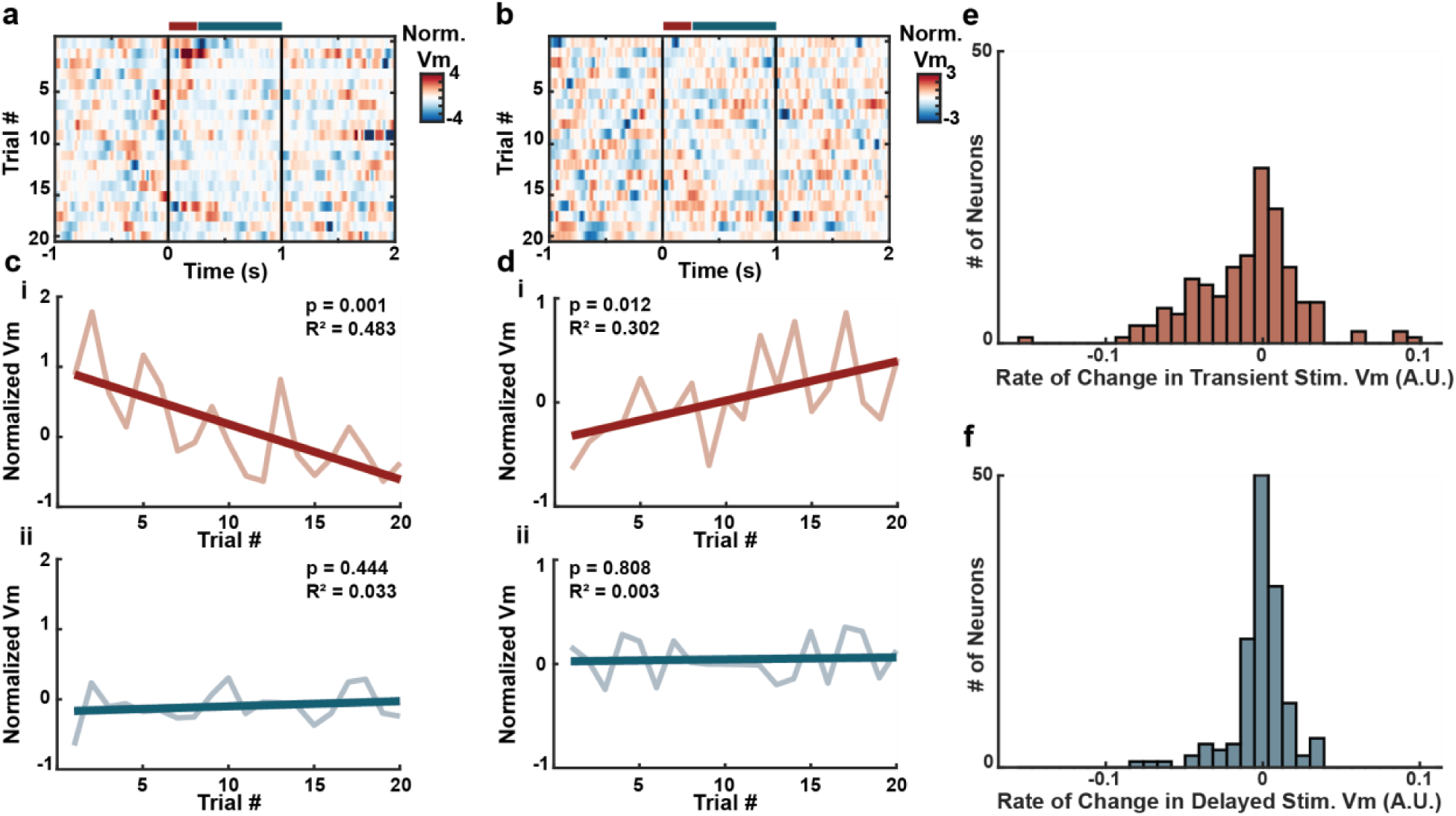
Evoked Vm responses across TUS trials. **(a–b)** Heatmaps of baseline-normalized Vm across trials for two example neurons showing progressive reduction **(a)** or increases **(b)** of evoked transient responses across trials. Each row represents one trial. Colored bars indicate analysis windows: red = transient period (0–250 ms), blue = delayed period (250–1000 ms). **(c–d)** Trial-by-trial evoked Vm response amplitude during the transient **(ci, di)** and delayed **(cii, dii)** periods for the neurons shown in a and (b), respectively. The darker lines are linear fits with the corresponding R^2^ and p-values indicated. **(e)** Distributions of the linear fit slope values (rate of change across trials) of the transient Vm across all recorded neurons (N = 161). **(f)** Same as (e), but for the delayed Vm.

### TUS entrained cortical neuron Vm at both 10 and 40 Hz PRFs

Upon observing rapid Vm changes following TUS onset, we further evaluated the temporal precision of these responses relative to individual TUS pulses by assessing Vm entrainment (**Fig. 5a,i**). Specifically, we computed the phase-locking value (PLV) between TUS pulse onsets and Vm, and comparing observed PLVs to a shuffled null distribution (α = 0.05, see **Methods**). We identified 22/106 (20.8%) significantly entrained neurons at 10 Hz and 7/55 (12.7%) at 40 Hz PRF (**Fig. 5c–d,k–l**). Sham control recordings showed chance-level entrainment at 5% (3/60), as expected with the expected false-positive rate. Thus, TUS delivered at physiologically relevant 10 and 40 Hz PRFs entrained many cortical neurons, consistent with prior findings that tonically firing Purkinje cells in anesthetized rats exhibited more temporally regular spiking in response to 50 and 100 Hz PRF TUS^84^.

**Figure 5:**
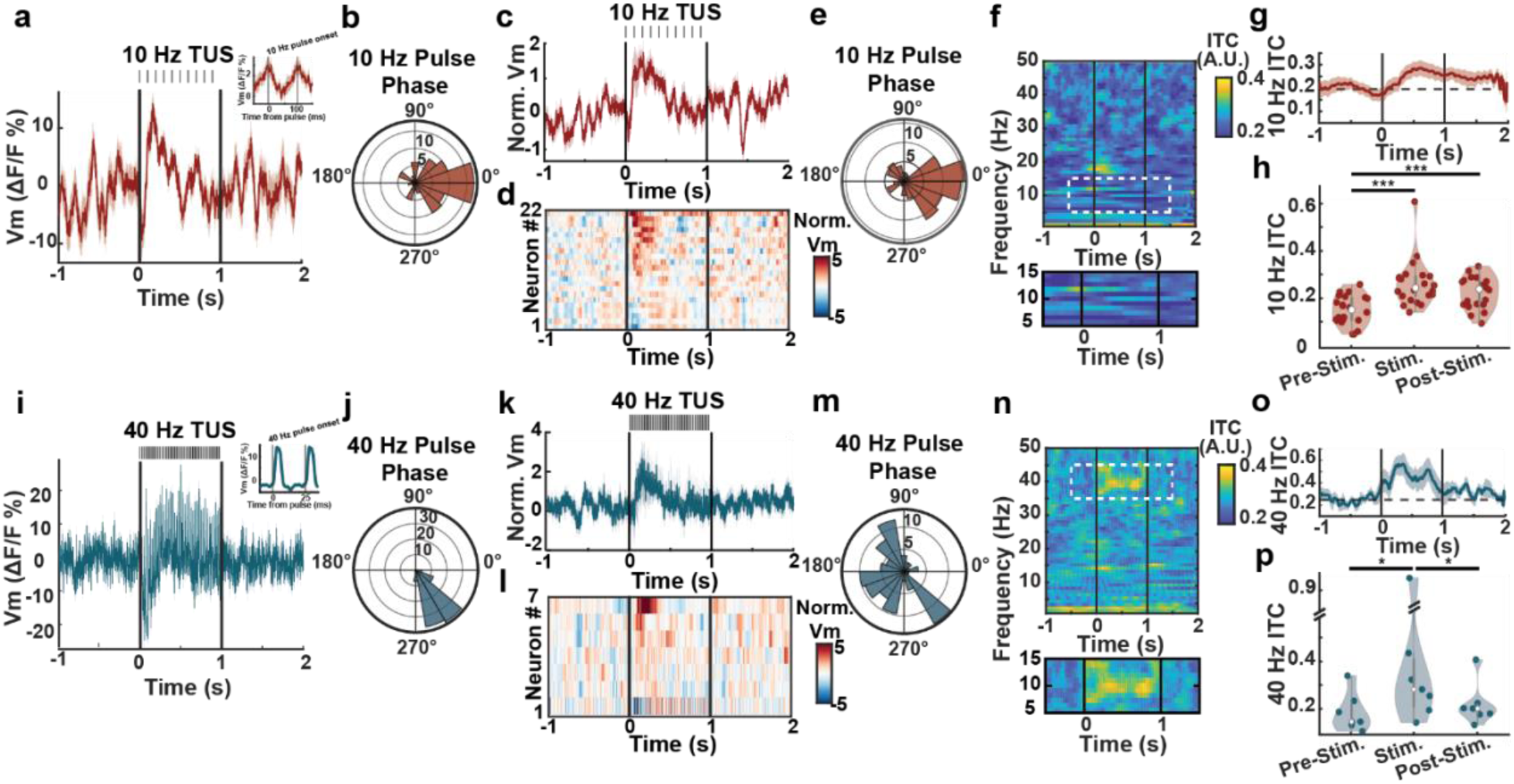
10 and 40 Hz TUS strongly entrains Vm and enhances inter-trial phase coherence. **(a)** Vm trace (ΔF/F%) from an example 10 Hz-entrained neuron with pulse-triggered average shown in the inset. **(b)** Polar plot of phase angles at pulse onset for the neuron shown in (a). **(c)** Z-scored Vm and **(d)** heatmap of all 10 Hz-entrained neurons (N = 22), sorted by response amplitude. **(e)** Circular distribution of pulse-onset phase angles across 10 Hz entrained neurons. **(f)** Inter-trial coherence (ITC) across frequencies aligned to stimulation onset. **(g)** ITC amplitude at 10 Hz. **(h)** Violin plots comparing ITC across pre-, during (stimulation), and post-TUS periods (Wilcoxon signed-rank test; see Suppl. Table 3). **(i–p)** Same as **(a–h)** for 40 Hz-entrained neurons (N = 7).

Vm fluctuations were tightly time-locked to stimulation pulses at both PRFs (**Fig. 5b,j**). For example, one 10 Hz-entrained neuron showed peak depolarization at 9 ms after each pulse, corresponding to a preferred phase angle of 2° (**Fig. 5b**), while a 40 Hz-entrained neuron showed a depolarization peak at 3 ms post-pulse, corresponding to a preferred phase angle of 303° (**Fig. 5j**). Across all 10 Hz entrained neurons, preferred phase angles clustered around 352±62.6° (mean±standard deviation, 22 neurons, N = 220 pulses), significantly deviated from uniformity (Rayleigh test; p = 1.28e-14, r = 0.40, N = 220 pulses, **Fig. 5e**). In contrast, 40 Hz-entrained neurons exhibited a broader distribution centered at 201±68.1° (mean±standard deviation, 7 neurons, N = 280 pulses), nonetheless significantly deviated from uniformity (Rayleigh test; p = 5.22e-9, r = 0.29, N = 280 pulses, **Fig. 5m**), indicating greater variability in response timing at 40 Hz PRF.

To further assess entrainment, we measured trial-to-trial phase consistency through inter-trial coherence (ITC) (**Fig. 5f,n, Methods**). ITC values were significantly different across pre-, during, and post-stimulation windows at both PRFs (**Fig. 5g,o**; Friedman tests; statistical results in **Suppl. Table 3**). ITC at both stimulation frequencies increased during stimulation relative to pre-stimulation, indicating phase-aligned responses across trials (Wilcoxon sign-rank tests; statistical results in **Suppl. Table 3**). At 10 Hz, ITC persisted significantly into the post-stimulation period, whereas 40 Hz, ITC returned to pre-stimulation levels (**Fig. 5h,p**). Together, these results demonstrate that TUS evoked precisely timed Vm responses at both 10 and 40 Hz PRFs, with 10 Hz PRF producing more consistent responses across individual pulses, highlighting the potential for frequency-specific pulsing protocols to modulate network dynamics through phase-dependent mechanisms.

### TUS resets network dynamics

As multiple neurons were recorded simultaneously (**Fig. 6a**), we further examined how TUS influences network dynamics by evaluating coordination between neurons. In many recordings, we noted that Vm and spiking became more coordinated immediately after TUS offset (**Fig. 6b**). To quantify this, we computed cross-correlations of Vm and spiking activity across simultaneously recorded neuron pairs (**Methods**). Both Vm and spike correlations varied significantly across conditions (Friedman test; statistical results in **Suppl. Table 3**), indicating that TUS alters inter-neuron coordination. Interestingly, correlations increased significantly post-stimulation compared to during TUS (Wilcoxon signed-rank test; see **Suppl. Table 3**), suggesting that TUS offset triggers a reset in network dynamics (**Fig. 6c,d**). This rebound synchrony may contribute to the plasticity effects observed (**Fig. 4**) and could be leveraged in therapeutic applications to modulate network states more effectively.

**Figure 6:**
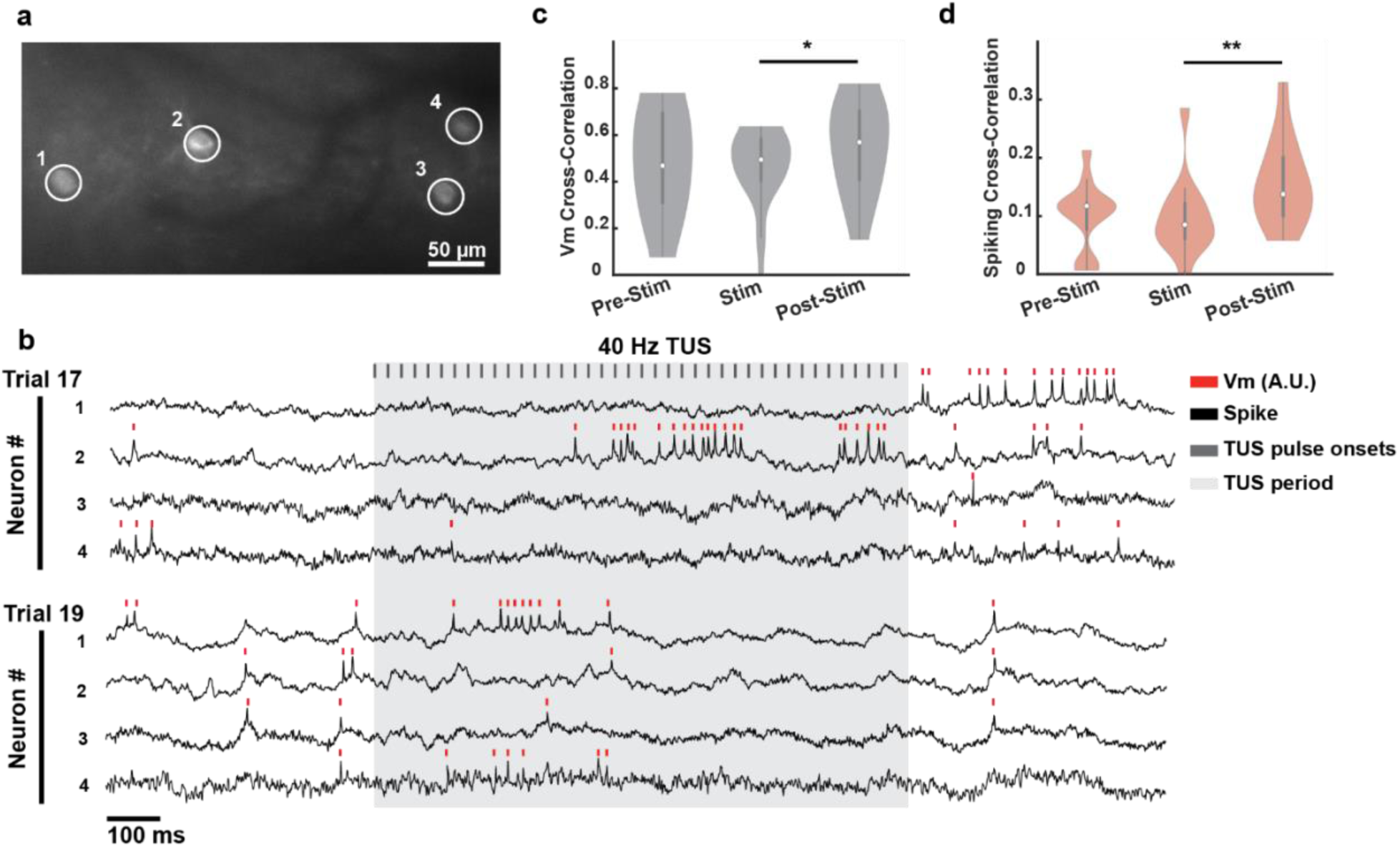
TUS modulates Vm and spiking synchronization between simultaneously recorded neurons. **(a)** SomArchon fluorescence from four simultaneously recorded neurons during 40 Hz TUS (scale bar: 50 µm). **(b)** Example Vm traces and spike times across two trials. Red ticks: spikes; black: Vm; gray ticks: pulse onsets; shaded gray: 1 s TUS (scale bar: 100 ms). **(c)** Violin plot of Vm zero-lag cross-correlations across pre-, during (stim), and post-TUS periods (N = 13 FOVs, Wilcoxon signed-rank test; see Suppl. Table 3). **(d)** Violin plot of zero-lag spike cross-correlations (N = 13 FOVs, Wilcoxon signed-rank test; see Suppl. Table 3).

## Discussion

Despite its clinical promise, the direct effect of TUS on individual neurons in the awake mammalian brain remains unknown. We employed high-speed cellular voltage imaging in awake mice to assess single-neuron Vm responses to TUS delivered at 10 and 40 Hz PRFs, free of TUS-mediated electromechanical artifacts often associated with electrodes^85^. We found that TUS robustly modulates individual neurons, producing diverse Vm changes, including depolarization, hyperpolarization, increased spiking, and entrainment. While 42.8% of recorded neurons showed significant Vm modulation, predominantly within 250 ms of stimulation onset, only 20.5% showed increased spiking, typically emerging after 250 ms, consistent with a gradual ramping depolarization evoked by TUS. Notably, many activated neurons responded with a latency shorter than 10 ms, suggesting direct neuronal effects. Reflecting the observed short-latency responses, many neurons were entrained by TUS at both frequencies (20.8% at 10 Hz; 12.7% at 40 Hz), with Vm fluctuations precisely time-locked to each pulse, especially during 10 Hz TUS, highlighting TUS as a temporally precise and biophysically selective neuromodulatory tool. Additionally, we discovered that the evoked Vm response magnitude systematically changed across many neurons, with most declining across repeated TUS trials, highlighting a plasticity mechanism during repeated TUS. In agreement with the plasticity effects, TUS reset network synchrony upon stimulation offset. These results demonstrate that pulsed TUS at physiologically relevant PRFs can directly drive temporally precise Vm changes in individual neurons and engage cellular plasticity processes supporting prolonged network modulation.

We observed rapid Vm changes with a latency shorter than 10 ms in many neurons, absent in sham recordings. Although transient onset and offset transitions of non-ramped TUS create broad frequency activation of the auditory brainstem within ∼6 ms^66^, propagation of cochlear activation requires 15–20ms to reach the auditory cortex^86^, which is beyond the latencies observed in our recordings. Thus, the short latency activation supports a direct and non-auditory origin of the evoked responses. Prior studies that observed ultrasound-evoked auditory responses used high PRFs of 1–1.5 kHz^63,64^, which are within the mouse hearing range (∼1–100 kHz^87,88^). We used a 0.35 MHz carrier frequency (well above auditory thresholds) and low PRFs (10 or 40 Hz) to minimize cochlear activation. The estimated peak pressure at the intracranial bone was 0.18 MPa, below the 0.22 MPa threshold previously modeled for auditory brainstem activation^65^. Furthermore, motor cortex neurons are minimally modulated by auditory stimuli without significant motor implications, further arguing against an auditory origin. Finally, off-target sham ultrasound evoked no significant changes in Vm and spiking, further ruling out non-specific systemic and auditory confounds. Thus, our results demonstrate a direct effect of TUS on neurons, which cannot be explained by auditory activation.

TUS evoked both excitatory and inhibitory Vm responses across cortical neurons, especially during the transient period within 250 ms of stimulation onset. Thus, restricting each TUS burst to 250 ms may optimize neural modulation while minimizing the overall acoustic energy delivery into the brain. Under the tested PRFs (10 and 40 Hz), response proportions and diversity were similar across groups, suggesting that cortical neurons exhibit broadly distributed sensitivity to these stimulation parameters. Prior work suggests that PRF can differentially modulate cell types, for instance, excitatory neurons increased firing with higher PRFs (range: 30 Hz to 4500 Hz), while inhibitory neurons remained largely unresponsive, even when duty cycle was held constant, suggesting intrinsic differences in temporal sensitivity^45^. Similarly, another study showed selective activation of PV+ interneurons population responses and suppression of CaMKII+ excitatory population responses in the hippocampus using 900 Hz continuous-wave stimulation, implicating both waveform shape and frequency as drivers of cell-type-specific divergence^61^. In the present study, low PRF (10 and 40 Hz) produced heterogeneous responses across cell types with no clear bias in evoked responses among CaMKIIα+ excitatory neurons, NDNF+ inhibitory interneurons, or a pan-neuronal Syn-expressing population (**Suppl. Table 4**). Thus, slower PRFs at 10 and 40 Hz broadly recruit cortical neurons with overlapping activation thresholds.

Beyond cell-type effects, accumulating evidence suggests that PRF alone can critically shape neural recruitment. In ex vivo slices, increasing PRF from 300 Hz to 1500 Hz markedly enhanced calcium response rates^40^. In humans, low-frequency TUS at 10–100 Hz produced sustained inhibition of motor-evoked potentials, while higher PRFs (1000 Hz) failed to elicit lasting effects, underscoring the nuanced influence of PRF on neuromodulatory outcomes^89^. Our findings extend this principle to the cellular level: although both 10 and 40 Hz stimulation reliably evoked rapid depolarizations, 40 Hz pulses produced slightly delayed response onset and earlier peak depolarization with a narrower temporal profile (reduced FWHM). These differences suggest that despite delivering less energy in the initial milliseconds, the higher temporal precision of 40 Hz stimulation may more efficiently trigger membrane responses through rapid pulse onsets and recovery intervals that minimize desensitization. In contrast, the longer 10 Hz pulse may yield more prolonged membrane engagement. These patterns suggest that PRF influences both the likelihood of activation and the temporal dynamics of Vm responses.

Across repeated trials, TUS-evoked neuronal responses within 250 ms of stimulation onset progressively shifted, with most neurons exhibiting response decay, but some showing augmentation, highlighting cellular adaptation. However, evoked responses during the delayed period, 250-1000 ms after stimulation onset, showed minimal changes over repeated TUS, consistent with the evoked Vm changes being primarily restricted to the first 250 ms of stimulation. This shift reflects adaptive plasticity mechanisms that regulate excitability during repeated stimulation. For example, activity-dependent recruitment of hyperpolarizing potassium conductances, which are upregulated by sustained depolarization or intracellular calcium^43,90^. TUS-mediated increase in intracellular calcium could thus contribute to the observed response depotentiation via recruitment of potassium channels. Another example is mechanosensitive ion channel Piezo1, which exhibits rapid inactivation under sustained or repetitive mechanical force. Piezo1 may enter desensitized states that reduce subsequent responsiveness to TUS^91^. Alternatively, repeated TUS may enhance inhibitory synaptic inputs via facilitation of GABAergic transmission or increased interneuron recruitment^92^. Synaptic depression at excitatory terminals, due to vesicle depletion or presynaptic calcium channel inactivation, may also contribute^93^. Finally, the polarity shift and response decay align with homeostatic plasticity mechanisms such as synaptic scaling or firing rate adaptation, which act to stabilize activity and prevent overexcitation^94,95^.

Beyond single-neuron Vm amplitude changes, our finding revealed that TUS can entrain cortical neurons to externally imposed frequencies. Significant phase-locking was observed in many neurons at both 10 and 40 Hz PRFs, with clear population-level clustering of preferred phase angles despite cell-to-cell variability. These results indicate that TUS can reliably bias the timing of membrane dynamics, aligning individual neurons to the temporal structure of stimulation. Prior mesoscopic studies have shown TUS-induced increases in LFP power at stimulation frequencies, including 40 Hz, which has been linked to enhanced memory and amyloid clearance in Alzheimer’s models^46,47^. Here, we extend these findings by resolving entrainment at the single-cell level in the cortex of awake mice. Compared to an earlier report in anesthetized rats, where tonically spiking Purkinje became more temporally regular during 50 and 100 Hz TUS^79^, our data demonstrates robust, phase-locked Vm entrainment across heterogeneous neocortical populations under more physiological conditions. The significant phase clustering suggests that TUS exerts a reliable mechanical drive across neurons, synchronizing Vm oscillations through stereotyped membrane displacements at each pulse.

EEG studies have reported increased signal complexity and entropy during TUS, reflecting decorrelated activity in broader cortical networks^96^. During TUS, we observed a slight reduction in cortical network synchrony, though not significant (**Suppl. Table 3**). However, we detected a statistically significant increase in synchronization between simultaneously recorded neurons for both Vm and spiking activity upon TUS offset. This network reset after TUS offset could contribute to the plasticity effects observed during repeated stimulation (**Fig. 4**). The rebound increase in synchrony is likely due to a subtle reduction during stimulation, which may not have achieved statistical significance due to limited sample size. Future studies with larger neuronal populations and longer recording durations may uncover subtle but functionally relevant changes.

## Acknowledgements

XH acknowledges funding from NSF (CBET-1848029 and CIF-1955981) and NIH R01NS109794. JS acknowledges NIH T32-GM008541. EB acknowledges NSF GRFP DGE-1840990 and NIH T32-EB006359. ES acknowledges NIH T32-GM008764 and NIH F31 NS143166-01. The authors additionally acknowledge support from the Shared Computing Cluster in Boston University’s Research Computing Services.

## Notes

### Competing Interest Statement

The authors have declared no competing interest.

## References

1. Fry FJ, Ades HW, Fry WJ. Production of Reversible Changes in the Central Nervous System by Ultrasound. Science. 1958;127(3289):83–84. doi:10.1126/science.127.3289.83

2. Tufail Y, Matyushov A, Baldwin N, et al. Transcranial Pulsed Ultrasound Stimulates Intact Brain Circuits. Neuron. 2010;66(5):681–694. doi:10.1016/j.neuron.2010.05.008

3. Ye PP, Brown JR, Pauly KB. Frequency dependence of ultrasound neurostimulation in the mouse brain. Ultrasound Med Biol. 2016;42(7):1512–1530. doi:10.1016/j.ultrasmedbio.2016.02.012

4. Ai L, Bansal P, Mueller JK, Legon W. Effects of transcranial focused ultrasound on human primary motor cortex using 7T fMRI: a pilot study. BMC Neurosci. 2018;19(1):56. doi:10.1186/s12868-018-0456-6

5. Kim T, Park C, Chhatbar PY, et al. Effect of Low Intensity Transcranial Ultrasound Stimulation on Neuromodulation in Animals and Humans: An Updated Systematic Review. Front Neurosci. 2021;15. Accessed April 21, 2022. https://www.frontiersin.org/article/10.3389/fnins.2021.620863

6. Colucci V, Strichartz G, Jolesz F, Vykhodtseva N, Hynynen K. Focused Ultrasound Effects on Nerve Action Potential *in vitro*. Ultrasound Med Biol. 2009;35(10):1737–1747. doi:10.1016/j.ultrasmedbio.2009.05.002

7. Kim H, Taghados SJ, Fischer K, Maeng LS, Park S, Yoo SS. Noninvasive Transcranial Stimulation of Rat Abducens Nerve by Focused Ultrasound. Ultrasound Med Biol. 2012;38(9):1568–1575. doi:10.1016/j.ultrasmedbio.2012.04.023

8. Downs ME, Lee SA, Yang G, Kim S, Wang Q, Konofagou EE. Non-invasive peripheral nerve stimulation via focused ultrasound in vivo. Phys Med Biol. 2018;63(3):035011. doi:10.1088/1361-6560/aa9fc2

9. Lin JW, Yu F, Müller WS, Ehnholm G, Okada Y. Focused ultrasound transiently increases membrane conductance in isolated crayfish axon. J Neurophysiol. 2019;121(2):480–489. doi:10.1152/jn.00541.2018

10. Yu F, Müller WS, Ehnholm G, Okada Y, Lin JW. Ultrasound-Induced Membrane Hyperpolarization in Motor Axons and Muscle Fibers of the Crayfish Neuromuscular Junction. Ultrasound Med Biol. 2023;49(12):2527–2536. doi:10.1016/j.ultrasmedbio.2023.08.016

11. Tyler WJ, Lani SW, Hwang GM. Ultrasonic modulation of neural circuit activity. Curr Opin Neurobiol. 2018;50:222–231. doi:10.1016/j.conb.2018.04.011

12. Kubanek J, Shukla P, Das A, Baccus SA, Goodman MB. Ultrasound Elicits Behavioral Responses through Mechanical Effects on Neurons and Ion Channels in a Simple Nervous System. J Neurosci. 2018;38(12):3081–3091. doi:10.1523/JNEUROSCI.1458-17.2018

13. Sorum B, Rietmeijer RA, Gopakumar K, Adesnik H, Brohawn SG. Ultrasound activates mechanosensitive TRAAK K+ channels through the lipid membrane. Proc Natl Acad Sci. 2021;118(6). doi:10.1073/pnas.2006980118

14. Yoo S, Mittelstein DR, Hurt RC, Lacroix J, Shapiro MG. Focused ultrasound excites cortical neurons via mechanosensitive calcium accumulation and ion channel amplification. Nat Commun. 2022;13(1):493. doi:10.1038/s41467-022-28040-1

15. Sherman J, Bortz E, Antonio ES, Tseng H an, Raiff L, Han X. Ultrasound pulse repetition frequency preferentially activates different neuron populations independent of cell type. Published online March 29, 2024:2024.03.25.586645. doi:10.1101/2024.03.25.586645

16. Qiu Z, Guo J, Kala S, et al. The Mechanosensitive Ion Channel Piezo1 Significantly Mediates In Vitro Ultrasonic Stimulation of Neurons. iScience. 2019;21:448–457. doi:10.1016/j.isci.2019.10.037

17. Hoffman BU, Baba Y, Lee SA, Tong CK, Konofagou EE, Lumpkin EA. Focused ultrasound excites action potentials in mammalian peripheral neurons in part through the mechanically gated ion channel PIEZO2. Proc Natl Acad Sci. 2022;119(21):e2115821119. doi:10.1073/pnas.2115821119

18. Clennell B, Steward TGJ, Hanman K, et al. Ultrasound modulates neuronal potassium currents via ionotropic glutamate receptors. Brain Stimulat. 2023;16(2):540–552. doi:10.1016/j.brs.2023.01.1674

19. Krasovitski B, Frenkel V, Shoham S, Kimmel E. Intramembrane cavitation as a unifying mechanism for ultrasound-induced bioeffects. Proc Natl Acad Sci. 2011;108(8):3258–3263. doi:10.1073/pnas.1015771108

20. Plaksin M, Kimmel E, Shoham S. Cell-Type-Selective Effects of Intramembrane Cavitation as a Unifying Theoretical Framework for Ultrasonic Neuromodulation. eNeuro. 2016;3(3):1–16. doi:10.1523/eneuro.0136-15.2016

21. Lemaire T, Neufeld E, Kuster N, Micera S. Understanding ultrasound neuromodulation using a computationally efficient and interpretable model of intramembrane cavitation. J Neural Eng. 2019;16(4):046007. doi:10.1088/1741-2552/ab1685

22. Tyler WJ, Tufail Y, Finsterwald M, Tauchmann ML, Olson EJ, Majestic C. Remote excitation of neuronal circuits using low-intensity, low-frequency ultrasound. PLoS ONE. 2008;3(10). doi:10.1371/journal.pone.0003511

23. Study Details | Investigating LFP Correlates of TUS in Patients With Movement Disorders | ClinicalTrials.gov. Accessed July 11, 2024. https://clinicaltrials.gov/study/NCT05965960?cond=Parkinson&intr=low%20intensity%20ultrasound&rank=6

24. Study Details | Effects of Peripheral Electrical Stimulation With Focused Ultrasound on Motor Symptoms in Parkinson’s Patients | ClinicalTrials.gov. Accessed July 11, 2024. https://clinicaltrials.gov/study/NCT06090292?cond=Parkinson&intr=low%20intensity%20ultrasound&rank=5

25. Study Details | Towards Noninvasive DBS of the Basal Ganglia in Parkinson’s Disease Using TUS | ClinicalTrials.gov. Accessed July 11, 2024. https://clinicaltrials.gov/study/NCT06232629?cond=Parkinson&intr=low%20intensity%20ultrasound&rank=4

26. Study Details | Non-invasive Neurostimulation in Parkinson’s Disease | ClinicalTrials.gov. Accessed July 11, 2024. https://clinicaltrials.gov/study/NCT01615718?cond=Parkinson&intr=low%20intensity%20ultrasound&rank=3

27. Study Details | LIFUP for Treatment of Motor Deficits in Parkinson’s Disease | ClinicalTrials.gov. Accessed July 11, 2024. https://clinicaltrials.gov/study/NCT04593875?cond=Parkinson&intr=low%20intensity%20ultrasound&rank=2

28. Study Details | Using Low Intensity Focused Ultrasound to Modulate Deep Brain Areas for Tremor Control in Parkinson’s Disease Patients. | ClinicalTrials.gov. Accessed June 26, 2024. https://clinicaltrials.gov/study/NCT06259708

29. Bratsos S, Karponis D, Saleh SN, Bratsos S, Karponis D, Saleh SN. Efficacy and Safety of Deep Brain Stimulation in the Treatment of Parkinson’s Disease: A Systematic Review and Meta-analysis of Randomized Controlled Trials. Cureus. 2018;10(10). doi:10.7759/cureus.3474

30. Vissani M, Isaias IU, Mazzoni A. Deep brain stimulation: a review of the open neural engineering challenges. J Neural Eng. 2020;17(5):051002. doi:10.1088/1741-2552/abb581

31. De Risio L, Borgi M, Pettorruso M, et al. Recovering from depression with repetitive transcranial magnetic stimulation (rTMS): a systematic review and meta-analysis of preclinical studies. Transl Psychiatry. 2020;10(1):1–19. doi:10.1038/s41398-020-01055-2

32. Bulteau S, Laurin A, Pere M, et al. Intermittent theta burst stimulation (iTBS) versus 10 Hz high-frequency repetitive transcranial magnetic stimulation (rTMS) to alleviate treatment-resistant unipolar depression: A randomized controlled trial (THETA-DEP). Brain Stimulat. 2022;15(3):870–880. doi:10.1016/j.brs.2022.05.011

33. Chung SW, Hoy KE, Fitzgerald PB. Theta-burst stimulation: a new form of TMS treatment for depression? Depress Anxiety. 2015;32(3):182–192. doi:10.1002/da.22335

34. Lan XJ, Yang XH, Qin ZJ, et al. Efficacy and safety of intermittent theta burst stimulation versus high-frequency repetitive transcranial magnetic stimulation for patients with treatment-resistant depression: a systematic review. Front Psychiatry. 2023;14. doi:10.3389/fpsyt.2023.1244289

35. Vogeti S, Boetzel C, Herrmann CS. Entrainment and Spike-Timing Dependent Plasticity – A Review of Proposed Mechanisms of Transcranial Alternating Current Stimulation. Front Syst Neurosci. 2022;16. doi:10.3389/fnsys.2022.827353

36. Booth SJ, Taylor JR, Brown LJE, Pobric G. The effects of transcranial alternating current stimulation on memory performance in healthy adults: A systematic review. Cortex. 2022;147:112–139. doi:10.1016/j.cortex.2021.12.001

37. Lowet E, Kondabolu K, Zhou S, et al. Deep brain stimulation creates informational lesion through membrane depolarization in mouse hippocampus. Nat Commun. 2022;13(1):7709. doi:10.1038/s41467-022-35314-1

38. Naor O, Krupa S, Shoham S. Ultrasonic neuromodulation. J Neural Eng. 2016;13(3):031003. doi:10.1088/1741-2560/13/3/031003

39. Burks SR, Lorsung RM, Nagle ME, Tu TW, Frank JA. Focused ultrasound activates voltage-gated calcium channels through depolarizing TRPC1 sodium currents in kidney and skeletal muscle. Theranostics. 2019;9(19):5517–5531. doi:10.7150/thno.33876

40. Manuel TJ, Kusunose J, Zhan X, et al. Ultrasound neuromodulation depends on pulse repetition frequency and can modulate inhibitory effects of TTX. Sci Rep. 2020;10(1):15347. doi:10.1038/s41598-020-72189-y

41. Berry MJ, Meister M. Refractoriness and Neural Precision. J Neurosci. 1998;18(6):2200-2211. doi:10.1523/JNEUROSCI.18-06-02200.1998

42. Sardi S, Vardi R, Tugendhaft Y, Sheinin A, Goldental A, Kanter I. Long anisotropic absolute refractory periods with rapid rise times to reliable responsiveness. Phys Rev E. 2022;105(1-1):014401. doi:10.1103/PhysRevE.105.014401

43. Bean BP. The action potential in mammalian central neurons. Nat Rev Neurosci. 2007;8(6):451–465. doi:10.1038/nrn2148

44. Ramachandran S, Gao H, Yttri E, Yu K, He B. Parameter-dependent cell-type specific effects of transcranial focused ultrasound stimulation in an awake head-fixed rodent model. J Neural Eng. 2025;22(2):026022. doi:10.1088/1741-2552/adbb1f

45. Yu K, Niu X, Krook-Magnuson E, He B. Intrinsic functional neuron-type selectivity of transcranial focused ultrasound neuromodulation. Nat Commun. 2021;12(1):2519. doi:10.1038/s41467-021-22743-7

46. Park M, Hoang GM, Nguyen T, et al. Effects of transcranial ultrasound stimulation pulsed at 40 Hz on Aβ plaques and brain rhythms in 5×FAD mice. Transl Neurodegener. 2021;10(1):48. doi:10.1186/s40035-021-00274-x

47. Chen J, Wang X, Li X, Li X, Zhang Y, Yuan Y. Ultrasound-Induced Synchronized Neural Activities at 40 Hz and 200 Hz Entrained Corresponded Oscillations and Improve Alzheimer’s Disease Memory. CNS Neurosci Ther. 2025;31(4):e70351. doi:10.1111/cns.70351

48. Iaccarino HF, Singer AC, Martorell AJ, et al. Gamma frequency entrainment attenuates amyloid load and modifies microglia. Nature. 2016;540(7632):230–235. doi:10.1038/nature20587

49. Yaakub SN, White TA, Roberts J, et al. Transcranial focused ultrasound-mediated neurochemical and functional connectivity changes in deep cortical regions in humans. Nat Commun. 2023;14(1):5318. doi:10.1038/s41467-023-40998-0

50. Oghli YS, Grippe T, Arora T, Hoque T, Darmani G, Chen R. Mechanisms of theta burst transcranial ultrasound induced plasticity in the human motor cortex. Brain Stimul Basic Transl Clin Res Neuromodulation. 2023;16(4):1135–1143. doi:10.1016/j.brs.2023.07.056

51. Kim HJ, Phan TT, Lee K, et al. Long-lasting forms of plasticity through patterned ultrasound-induced brainwave entrainment. Sci Adv. 2024;10(8):eadk3198. doi:10.1126/sciadv.adk3198

52. Niu X, Yu K, He B. Transcranial focused ultrasound induces sustained synaptic plasticity in rat hippocampus. Brain Stimulat. 2022;15(2):352–359. doi:10.1016/j.brs.2022.01.015

53. Murphy KR, Farrell JS, Bendig J, et al. Optimized ultrasound neuromodulation for non-invasive control of behavior and physiology. Neuron. 2024;112(19):3252–3266.e5. doi:10.1016/j.neuron.2024.07.002

54. Di Ianni T, Morrison KP, Yu B, Murphy KR, de Lecea L, Airan RD. High-throughput ultrasound neuromodulation in awake and freely behaving rats. Brain Stimulat. 2023;16(6):1743–1752. doi:10.1016/j.brs.2023.11.014

55. Ai L, Mueller JK, Grant A, Eryaman Y, Legon W. Transcranial focused ultrasound for BOLD fMRI signal modulation in humans. In: 2016 38th Annual International Conference of the IEEE Engineering in Medicine and Biology Society (EMBC). ; 2016:1758–1761. doi:10.1109/EMBC.2016.7591057

56. Yang PF, Phipps MA, Jonathan S, et al. Bidirectional and state-dependent modulation of brain activity by transcranial focused ultrasound in non-human primates. Brain Stimulat. 2021;14(2):261–272. doi:10.1016/j.brs.2021.01.006

57. Wang Y, Xie P, Zhou S, Wang X, Yuan Y. Low-intensity pulsed ultrasound modulates multi-frequency band phase synchronization between LFPs and EMG in mice. J Neural Eng. 2019;16(2). doi:10.1088/1741-2552/ab0879

58. Wang X, Yan J, Wang Z, Li X, Yuan Y. Neuromodulation Effects of Ultrasound Stimulation Under Different Parameters on Mouse Motor Cortex. IEEE Trans Biomed Eng. 2020;67(1):291–297. doi:10.1109/TBME.2019.2912840

59. Tseng H an, Sherman J, Bortz E, et al. Region-Specific Effects of Ultrasound on Individual Neurons in the Awake Mammalian Brain. iScience. Published online August 8, 2021:102955. doi:10.1016/j.isci.2021.102955

60. Cheng Z, Wang C, Wei B, Gan W, Zhou Q, Cui M. High resolution ultrasonic neural modulation observed via in vivo two-photon calcium imaging. Brain Stimulat. 2022;15(1):190–196. doi:10.1016/j.brs.2021.12.005

61. Murphy KR, Farrell JS, Gomez JL, et al. A tool for monitoring cell type–specific focused ultrasound neuromodulation and control of chronic epilepsy. Proc Natl Acad Sci. 2022;119(46):e2206828119. doi:10.1073/pnas.2206828119

62. Oh SJ, Lee JM, Kim HB, et al. Ultrasonic Neuromodulation via Astrocytic TRPA1. Curr Biol. 2019;29(20):3386–3401.e8. doi:10.1016/j.cub.2019.08.021

63. Guo H, Hamilton M, Offutt SJ, et al. Ultrasound Produces Extensive Brain Activation via a Cochlear Pathway. Neuron. 2018;98(5):1020–1030.e4. doi:10.1016/j.neuron.2018.04.036

64. Sato T, Shapiro MG, Tsao DY. Ultrasonic Neuromodulation Causes Widespread Cortical Activation via an Indirect Auditory Mechanism. Neuron. 2018;98(5):1031–1041.e5. doi:10.1016/j.neuron.2018.05.009

65. Choi MH, Li N, Popelka G, Butts Pauly K. Development and validation of a computational method to predict unintended auditory brainstem response during transcranial ultrasound neuromodulation in mice. Brain Stimulat. 2023;16(5):1362–1370. doi:10.1016/j.brs.2023.09.004

66. Mohammadjavadi M, Ye PP, Xia A, Brown J, Popelka G, Pauly KB. Elimination of peripheral auditory pathway activation does not affect motor responses from ultrasound neuromodulation. Brain Stimulat. 2019;12(4):901–910. doi:10.1016/j.brs.2019.03.005

67. Wang Z, Yan J, Wang X, Yuan Y, Li X. Transcranial Ultrasound Stimulation Directly Influences the Cortical Excitability of the Motor Cortex in Parkinsonian Mice. Mov Disord. 2020;35(4):693–698. doi:10.1002/mds.27952

68. Vion-Bailly J, N’Djin WA, Suarez Castellanos IM, Mestas JL, Carpentier A, Chapelon JY. A causal study of the phenomenon of ultrasound neurostimulation applied to an in vivo invertebrate nervous model. Sci Rep. 2019;9(1):13738. doi:10.1038/s41598-019-50147-7

69. Ibsen S, Tong A, Schutt C, Esener S, Chalasani SH. Sonogenetics is a non-invasive approach to activating neurons in Caenorhabditis elegans. Nat Commun. 2015;6(1):8264. doi:10.1038/ncomms9264

70. Kubanek J, Shukla P, Das A, Baccus SA, Goodman MB. Ultrasound elicits behavioral responses through mechanical effects on neurons and ion channels in a simple nervous system. J Neurosci. 2018;38(12):3081–3091. doi:10.1523/JNEUROSCI.1458-17.2018

71. Weinreb E, Moses E. Mechanistic insights into ultrasonic neurostimulation of disconnected neurons using single short pulses. Brain Stimulat. 2022;15(3):769–779. doi:10.1016/j.brs.2022.05.004

72. Truong TT, Chiu WT, Lai YS, Huang H, Jiang X, Huang CC. Ca2+ signaling–mediated low-intensity pulsed ultrasound–induced proliferation and activation of motor neuron cells. Ultrasonics. 2022;124:106739. doi:10.1016/j.ultras.2022.106739

73. Piatkevich KD, Bensussen S, Tseng H an, et al. Population imaging of neural activity in awake behaving mice. Nature. 2019;574(7778):413–417. doi:10.1038/s41586-019-1641-1

74. Sohal VS, Zhang F, Yizhar O, Deisseroth K. Parvalbumin neurons and gamma rhythms enhance cortical circuit performance. Nature. 2009;459(7247):698–702. doi:10.1038/nature07991

75. Martinez-Losa M, Tracy TE, Ma K, et al. Nav1.1-Overexpressing Interneuron Transplants Restore Brain Rhythms and Cognition in a Mouse Model of Alzheimer’s Disease. Neuron. 2018;98(1):75–89.e5. doi:10.1016/j.neuron.2018.02.029

76. Boraud T, Brown P, Goldberg JA, Graybiel AM, Magill PJ. Oscillations in the Basal Ganglia: The good, the bad, and the unexpected. In: Bolam JP, Ingham CA, Magill PJ, eds. The Basal Ganglia VIII. Advances in Behavioral Biology. Springer US; 2005:1–24. doi:10.1007/0-387-28066-9_1

77. Xiao S, Cunningham W, Kondabolu K, et al. Large-scale deep tissue voltage imaging with targeted-illumination confocal microscopy. Nat Methods. 2024;21. doi:10.1038/s41592-024-02275-w

78. Kim HJ, Phan TT, Lee K, et al. Long-lasting forms of plasticity through patterned ultrasound-induced brainwave entrainment. Sci Adv. 2024;10(8):eadk3198. doi:10.1126/sciadv.adk3198

79. Asan AS, Kang Q, Oralkan Ö, Sahin M. Entrainment of cerebellar Purkinje cell spiking activity using pulsed ultrasound stimulation. Brain Stimulat. 2021;14(3):598–606. doi:10.1016/j.brs.2021.03.004

80. Wattiez N, Constans C, Deffieux T, et al. Transcranial ultrasonic stimulation modulates single-neuron discharge in macaques performing an antisaccade task. Brain Stimulat. 2017;10(6):1024–1031. doi:10.1016/j.brs.2017.07.007

81. Clennell B, Steward TGJ, Elley M, et al. Transient ultrasound stimulation has lasting effects on neuronal excitability. Brain Stimulat. 2021;14(2):217–225. doi:10.1016/j.brs.2021.01.003

82. Grippe T, Shamli-Oghli Y, Darmani G, et al. Plasticity-Induced Effects of Theta Burst Transcranial Ultrasound Stimulation in Parkinson’s Disease. Mov Disord Off J Mov Disord Soc. Published online May 24, 2024. doi:10.1002/mds.29836

83. Folloni D, Verhagen L, Mars RB, et al. Manipulation of Subcortical and Deep Cortical Activity in the Primate Brain Using Transcranial Focused Ultrasound Stimulation. Neuron. 2019;101(6):1109–1116.e5. doi:10.1016/j.neuron.2019.01.019

84. Asan AS, Lang EJ, Sahin M. Entrainment of cerebellar purkinje cells with directional AC electric fields in anesthetized rats. Brain Stimulat. 2020;13(6):1548–1558. doi:10.1016/j.brs.2020.08.017

85. Collins MN, Mesce KA. Focused Ultrasound Neuromodulation and the Confounds of Intracellular Electrophysiological Investigation. eNeuro. 2020;7(4):ENEURO.0213–20.2020. doi:10.1523/ENEURO.0213-20.2020

86. James NM, Gritton HJ, Kopell N, Sen K, Han X. Muscarinic receptors regulate auditory and prefrontal cortical communication during auditory processing. Neuropharmacology. 2019;144:155–171. doi:10.1016/j.neuropharm.2018.10.027

87. Heffner HE, Heffner RS. Hearing ranges of laboratory animals. J Am Assoc Lab Anim Sci JAALAS. 2007;46(1):20–22.

88. Koay G, Heffner RS, Heffner HE. Behavioral audiograms of homozygous medJ mutant mice with sodium channel deficiency and unaffected controls. Hear Res. 2002;171(1):111–118. doi:10.1016/S0378-5955(02)00492-6

89. Zadeh AK, Raghuram H, Shrestha S, et al. The effect of transcranial ultrasound pulse repetition frequency on sustained inhibition in the human primary motor cortex: A double-blind, sham-controlled study. Brain Stimul Basic Transl Clin Res Neuromodulation. 2024;17(2):476–484. doi:10.1016/j.brs.2024.04.005

90. Wang J jing, Li Y. KCNQ potassium channels in sensory system and neural circuits. Acta Pharmacol Sin. 2016;37(1):25–33. doi:10.1038/aps.2015.131

91. J W, M Y, Ah L, An M, B K, J G. Inactivation of Mechanically Activated Piezo1 Ion Channels Is Determined by the C-Terminal Extracellular Domain and the Inner Pore Helix. Cell Rep. 2017;21(9). doi:10.1016/j.celrep.2017.10.120

92. Isaacson JS, Scanziani M. How inhibition shapes cortical activity. Neuron. 2011;72(2):231–243. doi:10.1016/j.neuron.2011.09.027

93. Zucker RS, Regehr WG. Short-term synaptic plasticity. Annu Rev Physiol. 2002;64:355-405. doi:10.1146/annurev.physiol.64.092501.114547

94. Marder E, Goaillard JM. Variability, compensation and homeostasis in neuron and network function. Nat Rev Neurosci. 2006;7(7):563–574. doi:10.1038/nrn1949

95. Turrigiano G. Homeostatic Synaptic Plasticity: Local and Global Mechanisms for Stabilizing Neuronal Function. Cold Spring Harb Perspect Biol. 2012;4(1):a005736. doi:10.1101/cshperspect.a005736

96. Li X, Yang H, Yan J, Wang X, Li X, Yuan Y. Low-Intensity Pulsed Ultrasound Stimulation Modulates the Nonlinear Dynamics of Local Field Potentials in Temporal Lobe Epilepsy. Front Neurosci. 2019;13:287. doi:10.3389/fnins.2019.00287

97. Xiao S, Cunningham WJ, Kondabolu K, et al. Large-scale deep tissue voltage imaging with targeted illumination confocal microscopy. Published online July 21, 2023:2023.07.21.548930. doi:10.1101/2023.07.21.548930

98. Clinard S, Wettstone E, Moore D, Snell J, Padilla F, Eames M. Low-Cost 3-D Hydrophone Scanning Tank with MATLAB GUI Control. Ultrasound Med Biol. 2022;48(1):157–163. doi:10.1016/j.ultrasmedbio.2021.09.022

99. E C, N K, Sk H, Rm H, Jt H. Development of a high resolution three-dimensional surgical atlas of the murine head for strains 129S1/SvImJ and C57Bl/6J using magnetic resonance imaging and micro-computed tomography. Neuroscience. 2007;144(2). doi:10.1016/j.neuroscience.2006.08.080

100. Pnevmatikakis EA, Giovannucci A. NoRMCorre: An online algorithm for piecewise rigid motion correction of calcium imaging data. J Neurosci Methods. 2017;291:83–94. doi:10.1016/j.jneumeth.2017.07.031

101. Xiao S, Lowet E, Gritton HJ, et al. Large-scale voltage imaging in behaving mice using targeted illumination. iScience. 2021;24(11):103263. doi:10.1016/j.isci.2021.103263

