## Supplemental Material for "Transcranial ultrasound stimulation modulates neuronal membrane potentials across broad timescales in the awake mammalian brain"

**Supplemental Table 1: Experimental parameters for pulsed TUS protocols.**

| Pulse Repetition Frequency | Pulse Type | Pulse Duration | Ramp Duration | Ramp Shape | Repetition Interval |
|---|---|---|---|---|---|
| **10 Hz** | Pulse | 20.00 ms | 0 s | rectangular | 100.00 ms |
| | Pulse train | 1 s | 0 s | rectangular | |
| **40 Hz** | Pulse | 5.00 ms | 0 s | rectangular | 25.00 ms |
| | Pulse train | 1 s | 0 s | rectangular | |

**Supplemental Table 2: Free-field and in situ acoustic parameters for 0.35 MHz transducer.**

| ***Free field*** **$P_{SP}$** | ***in situ*** **$P_{SP}$** | ***in situ*** **$I_{SPPA}$** | ***in situ*** **$I_{SPTA}$** | **# cycles** | **TIC** | **MI** |
|---|---|---|---|---|---|---|
| 401 kPa | 107 kPa | 0.129 W-cm$^{-2}$ | 25.8 mW-cm$^{-2}$ | 7,000 | 0.184 | 0.181 |

PSP = Spatial Peak Pressure; $I_{SPPA}$ = Spatial Peak Pulse Average Intensity; $I_{SPTA}$ = Spatial Peak Temporal Average Intensity; TIC = Thermal Index in Cranial bone; MI = Mechanical Index. Free-field values were measured in water with a calibrated hydrophone; in situ values were estimated via k-Wave modeling. # cycles indicate total ultrasound cycles per trial.

**Supplemental Table 3: Statistical tests.**

| Figure | Description | Test | Comparison | p-value | N |
|---|---|---|---|---|---|
| Figure 3 | Vm 10 Hz | Wilcoxon signed-rank | Transient (+) vs pre-stim.; Delayed vs pre-stim.; Transient (-) vs pre-stim. | p = 1.17e-06; p = 0.004; p = 0.031 | N = 31; N = 6; N = 9 |
| | Vm 40 Hz | Wilcoxon signed-rank | Transient (+) vs pre-stim.; Delayed vs pre-stim.; Transient (-) vs pre-stim. | p = 4.38e-04; p = 0.500; p = 0.063 | N = 16; N = 2; N = 5 |
| | Spike rate 10 Hz | Wilcoxon signed-rank | Transient (+) vs pre-stim.; Delayed vs pre-stim.; | p = 0.016; p = 0.039 | N = 7; N = 9 |
| | Spike rate 40 Hz | Wilcoxon signed-rank | Transient (+) vs pre-stim.; Delayed vs pre-stim.; | p = 0.008; p = 0.359 | N = 8; N = 9 |
| Figure 5 | ITC 10 Hz | Friedman test | Pre-stim. vs stim. vs post-stim. | p = 1.14e-6 | N = 22 |
| | | Wilcoxon signed-rank | Pre-stim. vs stim.; Pre-stim. vs post-stim.; Stim vs post-stim. | p = 4.01e-5; p = 3.78e-4; p = 0.067 | |
| | ITC 40 Hz | Friedman test | Pre-stim. vs stim. vs post-stim. | p = 0.050 | N = 7 |
| | | Wilcoxon signed-rank | Pre-stim. vs stim.; Pre-stim. vs post-stim.; Stim. vs post-stim. | p = 0.047 p = 0.016; p = 0.047 | |

| | | | | | |
|---|---|---|---|---|---|
| Figure 6 | Vm cross-correlation | Friedman test | Pre-stim. vs stim. vs post-stim. | p = 0.016 | N = 13 |
| | | Wilcoxon signed-rank | Pre-stim. vs stim.;<br>Pre-stim. vs post-stim.;<br>Stim. vs post-stim. | p = 0.147<br>p = 0.057<br>p = 0.017 | |
| | Spiking cross-correlation | Friedman test | Pre-stim. vs stim. vs post-stim. | p = 0.004 | N = 13 |
| | | Wilcoxon signed-rank | Pre-stim. vs stim.;<br>Pre-stim. vs post-stim.;<br>Stim. vs post-stim. | p = 0.685;<br>p = 0.094;<br>p = 0.001 | |

Inter-trial coherence (ITC); All tests are two-tailed.

**Supplemental Table 4: Vm and spike modulation across cell types and PRFs.**

| Cell Type / PRF | Metric | Total Neurons | Modulated | Total Modulated (%) |
|---|---|---|---|---|
| **CaMKII** | **Vm** | **50** | **16** | **32.0** |
| *10 Hz* | Vm | 29 | 10 | 34.5 |
| *40 Hz* | Vm | 21 | 6 | 28.6 |
| **NDNF** | **Vm** | **23** | **13** | **56.5** |
| *10 Hz* | Vm | 9 | 5 | 55.6 |
| *40 Hz* | Vm | 14 | 8 | 57.1 |
| **Synapsin** | **Vm** | **88** | **43** | **48.9** |
| *10 Hz* | Vm | 68 | 32 | 47.1 |
| *40 Hz* | Vm | 20 | 11 | 55.0 |
| **Total** | **Vm** | **161** | **69** | **42.9** |
| **CaMKII** | Spike | **50** | **8** | **16.0** |
| *10 Hz* | Spike | 29 | 1 | 3.4 |
| *40 Hz* | Spike | 21 | 7 | 33.3 |
| **NDNF** | **Spike** | **23** | **7** | **30.4** |
| *10 Hz* | Spike | 9 | 4 | 44.4 |
| *40 Hz* | Spike | 14 | 3 | 21.4 |
| **Synapsin** | **Spike** | **88** | **18** | **20.5** |
| *10 Hz* | Spike | 68 | 11 | 16.2 |
| *40 Hz* | Spike | 20 | 7 | 35.0 |
| **Total** | **Spike** | **161** | **33** | **20.5** |
